# Seipin forms a flexible cage at lipid droplet formation sites

**DOI:** 10.1101/2021.08.05.455270

**Authors:** Henning Arlt, Xuewu Sui, Brayden Folger, Carson Adams, Xiao Chen, Roman Remme, Fred A. Hamprecht, Frank DiMaio, Maofu Liao, Joel M. Goodman, Robert V. Farese, Tobias C. Walther

**Affiliations:** Department of Molecular Metabolism, Harvard T. H. Chan School of Public Health, Boston, MA, 02115 USA; Department of Cell Biology, Harvard Medical School, Boston, MA, 02115 USA; Howard Hughes Medical Institute, Boston, MA, 02115 USA; Department of Pharmacology, University of Texas Southwestern Medical School, Dallas, TX 75390-9041, USA; Department of Biochemistry and Institute of Protein Design, University of Washington, Seattle, WA, 98195 USA; Heidelberg Collaborative for Image Processing, Interdisciplinary Center for Scientific Computing, Heidelberg University, 69120 Heidelberg, Germany; Broad Institute of Harvard and MIT, Cambridge, MA, 02124 USA

## Abstract

Lipid droplets (LDs) form in the endoplasmic reticulum by phase separation of neutral lipids. This process is facilitated by the seipin protein complex, which consists of a ring of seipin monomers, with yet unclear function. Here, we report a structure of yeast seipin based on cryo-electron microscopy and structural modeling data. Seipin forms a decameric, cage-like structure with the lumenal domains forming a stable ring at the cage floor and transmembrane segments forming the cage sides and top. The transmembrane segments interact with adjacent monomers in two distinct, alternating conformations. These conformations result from changes in switch regions, located between the lumenal domains and the transmembrane segments, that are required for seipin function. Our data suggest a model for LD formation in which a closed seipin cage enables TG phase separation and subsequently switches to an open conformation to allow LD growth and budding.

## INTRODUCTION

Lipid droplets (LDs) are cellular organelles with a primary function of storing lipids for energy generation and membrane biogenesis^1,2^. They serve as hubs of lipid metabolism, platforms for virus assembly, and organizing centers of innate immunity^3–5^. Although cellular LD formation is an evolutionarily conserved, fundamental process, its mechanism is still poorly understood. At its essence, LD biogenesis is the formation of emulsified oil droplets, driven by phase separation of enzymatically synthesized neutral lipids, such as triglycerides (TGs), within the lipid bilayer of the ER^6,7^. LDs subsequently bud into the cytoplasm. LD assembly protein complexes (LDACs) ensure the fidelity of this process and determine where LDs form^8,9^.

A key component of the LDAC is the evolutionarily conserved ER membrane protein seipin, encoded by the *BSCL2* gene in humans^10^. The importance of seipin in LD formation is emphasized by the phenotypes associated with seipin deficiency. In seipin-deficient yeast cells, LDs form inefficiently with TG blisters accumulating in the ER^11^. Moreover, LDs in these cells have abnormal protein composition^12^, and unstable junctions with the ER^13^. Similarly, mammalian cells lacking seipin form many abnormally small LDs with altered protein composition, as well as giant LDs^14^. In humans, seipin deficiency results in either lipodystrophy, multiple organ problems, and neurological defects, depending on the mutation^9^. Seipin consists of an evolutionarily conserved ER lumenal domain and flanking transmembrane segments, and less conserved cytoplasmic N- and C-terminal regions with lengths that vary among species (Fig. S1a).

Seipin monomers form a ~150 Å-diameter toroid complex, consisting of 12 or 11 subunits in flies or humans, respectively^15,16^. Within the complex, each lumenal domain folds into an α/β-sandwich domain with resemblance to lipid binding domains^15,16^. This domain is reported to bind negatively charged phospholipids^16^. The lumenal domains form a ring of hydrophobic helices oriented toward the center of the toroid complex and are predicted to insert into the lumenal leaflet of the ER membrane^15,17,18^. In mammals, these helices are necessary for seipin’s interaction with the LDAC accessory protein LD assembly factor 1 (LDAF1)^19^, which may be an orthologue of yeast LD organization (Ldo) proteins^20,21^. In contrast to flies or humans, yeast seipin (Sei1) requires another ER protein, Ldb16, for LDAC function in LD biogenesis^12,22^. Ldb16 has a long hydrophobic stretch with at least one transmembrane segment, but its function is unclear.

Based on experimental evidence, we proposed that LDACs catalyze neutral lipid accumulation and phase separation of neutral lipids in the ER, generating a neutral lipid lens at the LDAC^15,19^. Formation of LDs at LDACs occurs at lower TG concentrations than without seipin^19^. This model is supported by molecular simulation experiments utilizing the lumenal domain structures that detect TG molecules binding and accumulating at seipin’s central hydrophobic helices^17,18^. Other models for seipin function include generating or transferring specific lipids to forming LDs^23,24^, or promoting calcium transport^25,26^

Nonetheless, insights into how seipin and LDACs ensure the fidelity of LD formation have been lacking. One limitation for determining seipin and LDAC function is that structural information and analyses have been restricted so far to seipin’s lumenal domain^15,16^. Yet, mutations in this region have relatively minor or variable effects on LD formation^15,16,17^, suggesting that crucial determinants of seipin function may lie outside of this domain.

To gain further insight into seipin function, here we combined cryo-electron microscopy (cryo-EM) with deep learning-guided protein structure prediction based on evolutionary couplings^27^ to generate a full-length structural model of seipin. Validating and testing these structural predictions provide a new model for how seipin functions in LDACs to catalyze LD formation.

## RESULTS

### Seipin’s Transmembrane Segments Are Crucial for Its Function in LD Formation

We hypothesized that seipin’s evolutionarily conserved transmembrane (TM) segments are required for LD biogenesis. To test this idea, we replaced either seipin’s N-terminal or both TM segments with transmembrane helices from a structurally unrelated, human ER protein, FIT2^28^ (Fig. S1b). The resulting chimeric Sei1_(TM-N-FIT2)_ and Sei1_(TM-NC-FIT2)_ proteins, which were GFP-tagged, localized in puncta to the ER, in a pattern similar to wildtype (WT) seipin (Fig. S1c). We also constructed stable lines in which chromosomal seipin was tagged with 13xmyc and expression driven by either the endogenous promoter or the strong *PGK1* promoter, which generally equalized otherwise low expression of mutants (Fig. S1d). To test for an effect on oligomer formation, we isolated membranes and examined detergent-solubilized complexes by size-exclusion chromatography. WT seipin-myc migrated in two peaks – a large complex of an apparent mass well above the 669 kDa marker (see below), and a peak at an elution volume corresponding to ~300 kDa, likely representing micelles containing non-oligomerized seipin. Both mutant constructs showed WT-like oligomers, but no small 300-kDa peaks (Fig. S1e). To determine if the mutant constructs could rescue function, we analyzed the size of LDs in the stable cell lines. WT cells contained multiple small (r < 400 nm), relatively uniform LDs, whereas *sei1Δ* cells typically had tight clusters of small or supersized LDs (r > 400 nm; Fig. S1f;^29,30^). Neither Sei1(TM-N-FIT2) nor Sei1(TM-NC-FIT2) constructs rescued the null LD phenotype (Fig. S1f-h). Furthermore, neither mutant rescued the growth phenotype of *sei1Δ* cells on media containing terbinafine, a squalene epoxidase inhibitor, which serves as an alternative functional assay for seipin (Fig. S1i;^22^). These findings indicate that the transmembrane segments are crucial for seipin function.

### Molecular Structure of Yeast Seipin

To better understand seipin function, and particularly, the role of its TM segments, we sought to generate a molecular structure for the entire seipin protein. To this end, we purified the yeast seipin Sei1-Ldb16 complex by affinity and size-exclusion chromatography from a yeast strain that overproduced both proteins (Fig. S2a,b). After cleavage of the 2xProteinA tag conjugated to Sei1 with TEV protease, the oligomeric Sei1-Ldb16 complex migrated at an elution volume corresponding to ~600 kDa.

Initial processing of negative stain and cryo-EM images of the purified complex yielded a toroid structure of 10 subunits (Fig. S2c-e). Analysis of this density map with C_10_ symmetry revealed a region corresponding to seipin ER-lumenal domains resolved to an overall resolution of ~3.4Å, but with only weak densities of the TM segments (Fig. S2e). The poor resolution of the TM segments might have been due to heterogeneity in the conformations of the TM helices. To explore this possibility, we further classified the cryo-EM particle images without applying symmetry after C_10_ symmetry expansion (Fig. S2e). This revealed that densities from each class of particles visually resembled C_5_ symmetry. We refined the class with the highest predicted resolution with C_5_ symmetry, which revealed a ~145-Å diameter complex with two alternating conformations of the TM segments that were invisible in the 3D reconstruction with C_10_ symmetry. We designated these alternate conformations A and B (Fig. 1). In this map, nearly all of the lumenal domain of seipin and most of the transmembrane regions are well resolved with an overall resolution of ~3.2 Å (Fig. 1a,b; S3a,b).

**Figure 1:**
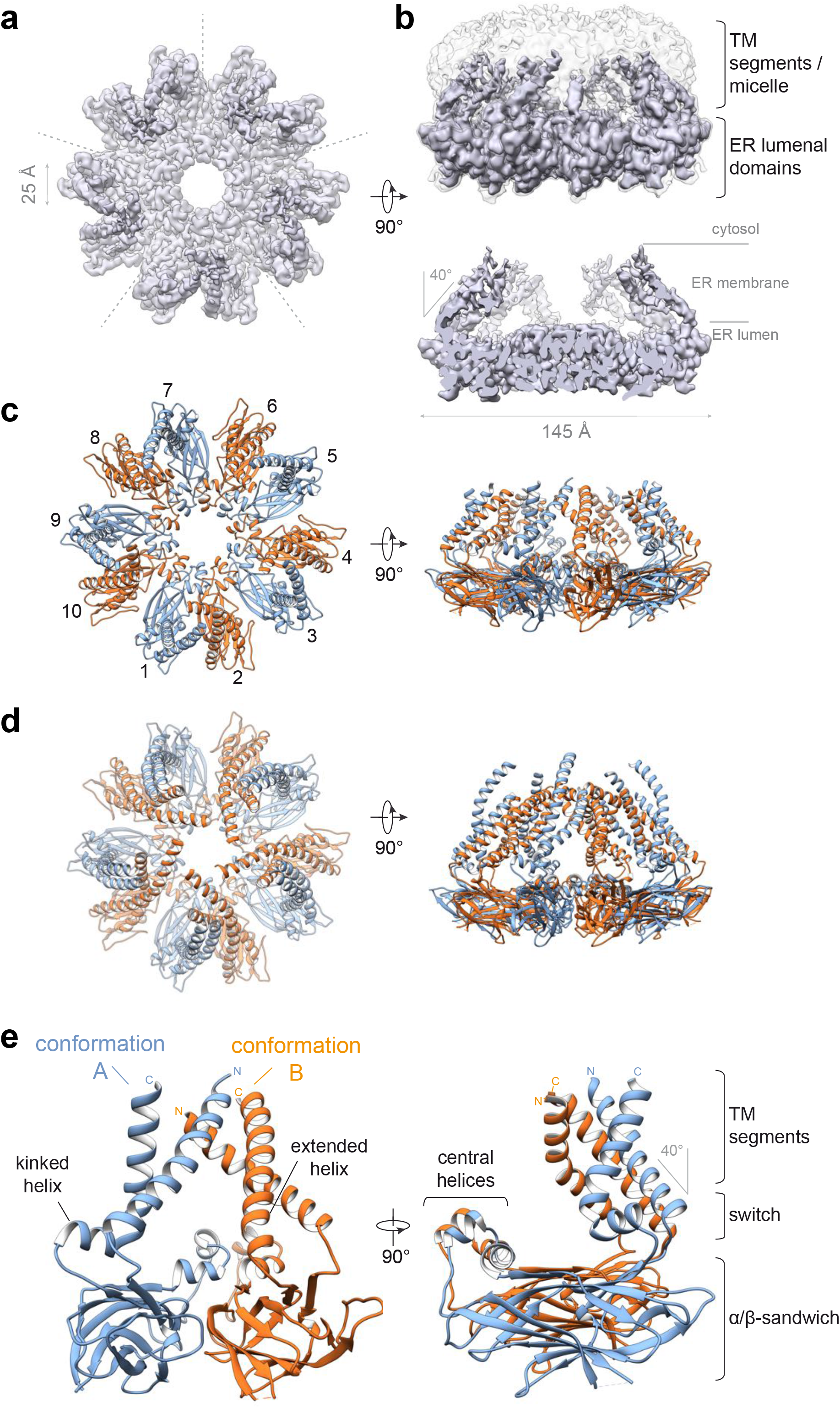
Cryo-EM structure of yeast seipin Sei1. **(a)** Cryo-EM density map of purified seipin oligomers shows the density of the lumenal domain and TM segments. The 5 symmetrical subunits are indicated by dashed lines. **(b)** Sideview of cryo-EM density map. Top, overlay of unsharpened density map (semi-transparent grey) showing the shape of the micelle, with sharpened map (purple). Bottom, sliced view of EM density map reveals cage-like structure. **(c,d)** Model of seipin show 10 seipin subunits per oligomer. Top view from the cytosolic side. (c) Model contains residues 17–264 for both A and B conformations, except loop residues 134– 147, which are not observed in the EM density map. (d) Extended structural model beyond EM density map contains residues 11–283 for conformation A (blue) and residues 8–285 for conformation B (orange) modelled by AI-assisted structure prediction. **(e)** Seipin oligomers contain two alternating monomer conformations termed A (blue) and B (orange) that differ only in the switch and TMD region, while the lumenal domains have the same structure.

Based on this EM density map, we built a molecular model of conformation A that included parts of both TM segments and the entire lumenal domain (amino acids (aa) 25–258), except a small segment of residues (aa 134–147) (Fig. S3c-e). The EM density for the TM segments of conformation B was of lower resolution than for conformation A (Fig. S3a,f), but nevertheless allowed us to manually build an initial model for the lumenal domain and connecting residues to the TM segments (residues 46–234). To build a model for the remainder of both TM segment conformations, we used Rosetta structural modelling, guided by both experimental electron density data, and distance and angle constraints generated by a deep neural network (trRosetta) trained to predict contacts from evolutionary couplings (Fig. S3g,h;^27^). This allowed placement of α-helices into the EM densities of conformation A (residues 17–25 and 258–264) and B (residues 17–45 and 235–264), producing a nearly complete model of the seipin protein backbone (Fig. 1c; S3f). This approach also allowed us to extend our model beyond what was resolved in the EM density map that contained almost all of the seipin sequence (conformation A, residues 11–283; conformation B, residues 8-285) (Fig. 1d). Although Ldb16 was detectable in the purified complex (Fig. S2b), all the protein density observed by cryo-EM could be unambiguously assigned to seipin.

Our model for yeast seipin revealed a decameric complex with the shape of a domed cage, with the lumenal domains forming the floor of the cage, predicted to sit beneath the lumenal leaflet of the ER membrane (Fig. 1b-d). All lumenal domains of the decameric complex had the same structure, with each lumenal domain containing an α/β-sandwich fold, similar to those in human and fly seipin^15,16^, and with two short central α-helices oriented toward the center ring of the cage floor (Fig. 1c-e). Two “switch” regions (residues 40–55 and 231–243), representing the biggest differences between conformations A and B, connect the ring of folded lumenal domains to the TM segments of seipin. The TM segments form the side walls of the cage and are tilted towards the center of the oligomer, coming together in a dome-shape at the cytoplasmic side of the complex. The architecture of a cage leads to a large, enclosed cavity in the center of the complex, predicted to be in the plane of the ER membrane (Fig. 1a-c).

In conformation A, the N-terminal transmembrane helix is tilted ~40° towards the center of the oligomer, whereas the C-terminal switch region adopts a kinked α-helix connected to the second TM helix (Fig. 1e). In conformation B, the C-terminal TM helix exhibits a continuously extended helix through the switch region and lacks the kink found in conformation A. As a result, the N-terminal TM helix in conformation B is tilted further (~60°) towards the center of the oligomeric assembly, and both TM segments lie close to the N-terminal TM helix of the neighboring conformation A monomer (Fig. 1c-e).

### Interactions in the Seipin Lumenal Domain Are Sufficient for Lumenal Domain Oligomerization but Are Not Required for Seipin Function

Comparing the architecture of individual seipin lumenal domains of yeast, fly, and human proteins revealed a striking difference (Figs. 2a,b). The fly and human lumenal domains possess a longer central helix that is hydrophobic, interacts with LDAF1 in humans^19^, and is implicated in binding TGs in molecular dynamics simulations^17,18^. In contrast, yeast seipin has two short helices with several charged residues (e.g., Q169, E172, Q173, E184) and a different orientation compared with human or fly seipin (Fig. 2a).

**Figure 2:**
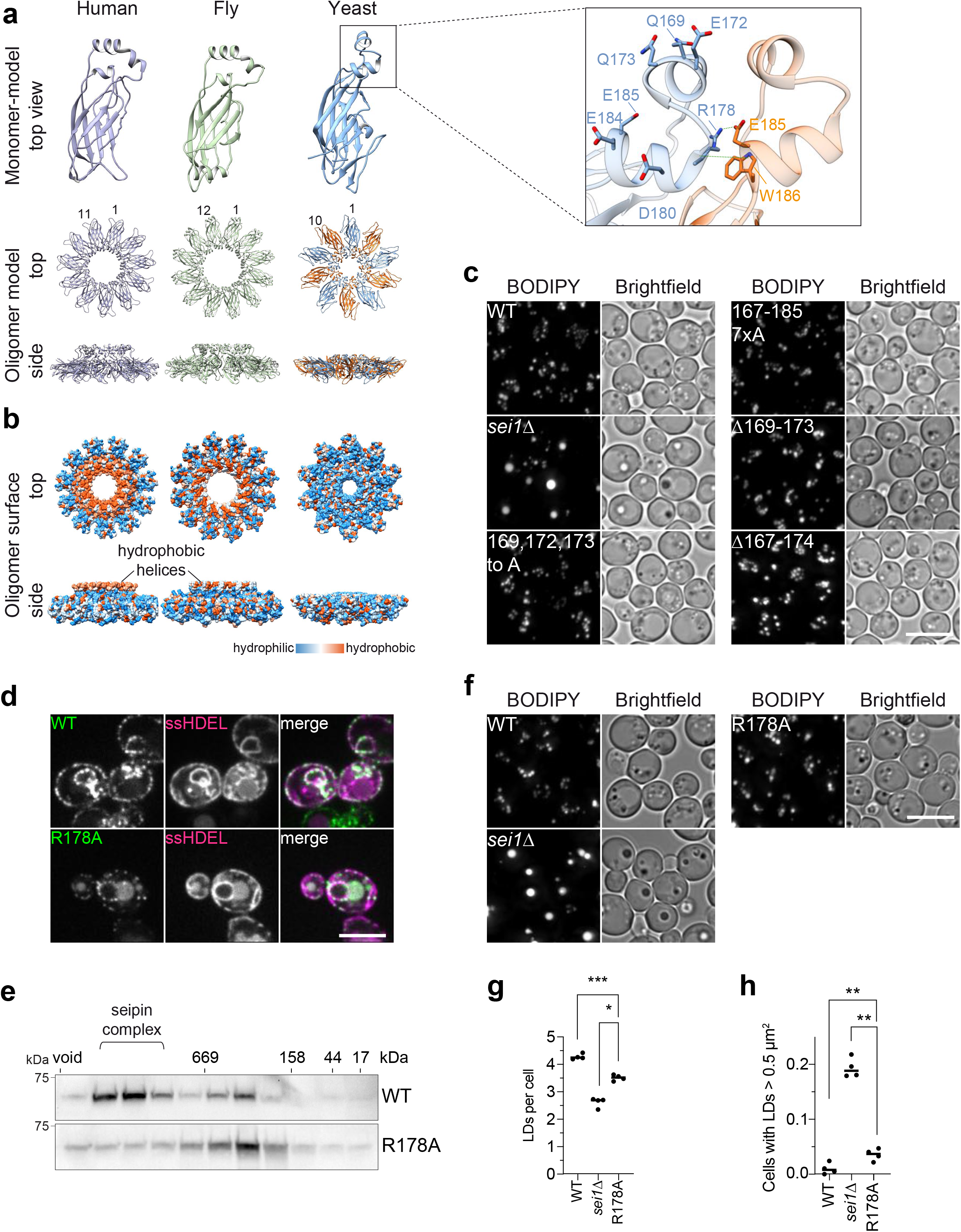
Interactions of seipin lumenal domain are sufficient for oligomerization but are not required for seipin function. **(a)** Comparison of seipin lumenal domain structural models of monomers and oligomers from fly (PDB 6MLU), human (PDB 6DS5), and yeast. Magnified box shows detailed view of yeast central helix, including neighboring monomer. **(b)** Hydrophobic surfaces of human, fly and yeast seipin lumenal domains indicate hydrophobic helices present in human and fly, but not yeast seipin. **(c)** LD morphology of strains expressing central helix mutants from seipin genomic locus. Cells were grown to high density and LDs were stained with BODIPY. **(d)** WT and R178A localize normally to the ER and form seipin foci. C-terminal GFP-tagged WT and R178A expressed from plasmids in *sei1Δ* cells. ssHDEL was also expressed from a plasmid. Size bar = 5 μm. **(e)** Seipin WT shows two peaks in size-exclusion chromatography of membrane extract in Triton X-100 from cell expressing *SEI1*-13xmyc WT from endogenous promoter or R178A mutant from *PGK1* promoter. Immunoblot with anti-myc antibodies. **(f)** Microscopy analysis of cell expressing indicated seipin mutants from endogenous locus driven by *PGK1* promoter with C-terminal 13xmyc tag or deleted for seipin (*sei1Δ*). Staining as in c. Size bar = 5 μm. **(g,h)** Quantification of LD morphology from the experiment shown in f. One dot indicates one separate experiment. *, p<0.05; **, p<0.01; ***, p<0.001.

To test whether residues in this central lumenal α-helix are important for yeast seipin function, we mutated residues Q169, E172, and Q173 or a combination of S167, Q169, E172, Q173, D180, E184, and E185 to alanine (169, 172, 173 to A; 167-185 7xA). Alternatively, we deleted the entire helical region and tested the functionality of these mutants in LD formation (Δ169-173 or Δ167-174). Cells expressing central lumenal α-helix mutations did so at normal or moderately reduced levels (Fig. S4a) and had LD phenotypes similar to WT (Fig. 2c; S4b,c). Additionally, each of these mutants complemented growth of seipin-deficient cells on media containing terbinafine (Fig. S4d). While *sei1Δ* cells had markedly decreased Ldb16 protein levels, mutants of the lumenal α-helix had normal or slightly decreased amounts of Ldb16, indicating this region is not required for binding and stabilization of Ldb16 by Sei1 (Fig. S4e)^22^.

Neighboring monomers of the lumenal domains appear to contact each other between residues R178 and E185/W186 of the adjacent monomers (Fig. 2a, inset) to form a hydrogen bond and a salt bridge between R178 and E185 (dotted green lines in Fig. 2a inset) and a cation-π interaction between R178 and W186. Because R178 is central to both interactions, we mutated this residue to alanine to determine if this interface is required for oligomer formation or stability. C-terminal GFP-tagged seipin R178A localized normally to the ER and formed characteristic GFP-puncta comparable in intensity to the WT protein (Fig. 2d), indicating normal oligomer formation *in vivo*. We integrated R178A containing a C-terminal 13x-myc tag into the endogenous seipin locus, which expressed at reduced levels (Fig. S5a), and examined oligomer stability in detergent extracts as described above. Unlike WT seipin that showed two peaks, the R178A mutant showed only the smaller ~300-kD peak, and this defect was not corrected by overexpression from the *PGK1* promoter (Fig. 2e; S5b), suggesting that R178A is important for decamer integrity, at least in detergent-solubilized seipin. A possible hypothesis for Ldb16 function is that it is an assembly factor for seipin complexes. However, deletion of *LDB16* had no effect on oligomerization of WT seipin, and overexpression of *LDB16* failed to rescue R178A oligomerization (Fig. S5b).

Seipin R178A only modestly affected LD morphology (Fig. 2f-h) and fully rescued the terbinafine sensitivity of *sei1Δ* cells (Fig. S5c). Mutation of other residues in the α/β-sandwich contact region (e.g., Q114A and E172A), alone or in combination with R178A, had no effect on LD phenotypes or terbinafine sensitivity in addition to R178A (Fig. S5d-f).

We further tested if seipin lumenal domains are sufficient for decamer formation by expressing a truncated version of the protein lacking transmembrane segments in *E. coli* (WT47-235) (Fig. S5g-i). By size-exclusion chromatography, the expressed lumenal domain (isolated in the absence of detergent) was sufficient to form oligomers and showed typical ring-shaped decameric assemblies visualized by negative staining electron microscopy (Fig. S5h,i). Introducing the R178A mutation into the isolated lumenal domains abrogated oligomerization (Fig. S5h), indicating that R178 is crucial for assembly of the decamer in the absence of the TM segments and may also be important for stability of the entire protein.

### Intramolecular Interactions of Transmembrane Helices Are Important for Seipin Function and Oligomer Formation

Within a monomer, both TM segments show close contacts (Fig. 3a). The significance of this is supported by an extensive network of trRosetta-predicted interactions between the N- and C-terminal TM segments (Fig. 3a-c). In particular, two patches of residues co-evolved and are predicted to interact within the monomer (e.g., residues S33-I259; Y37-Y248; Y41-M240; Fig. 3a-c). To test the requirement for these apparent evolutionary couplings, we mutated specific residues in the N-terminal transmembrane segment (Patch 1, S33A, Y37A, Y41A) or C-terminal transmembrane segment (Patch 2, M240G, Y248I, F255R, I259K). Mutating these patches did not affect seipin localization to the ER, although expression levels were lower than WT, and were restored by inserting the *PGK1* promoter (Fig. 3d,e; S6a). Analysis of oligomerization of the patch mutants by size-exclusion chromatography showed WT-like oligomers. However, combinations with the oligomerization mutant R178A led to unfolding or aggregation of the chimeras indicating higher instability of these mutants (Fig. 3f; S6b), and suggesting that the seipin transmembrane helices normally aids in decamer stability, which becomes critical in the absence of R178 lumenal interactions. Expression of mutants in patch 2, or patches 1 and 2 in *sei1Δ* cells did not maintain seipin function in LD morphology or growth on terbinafine-containing medium, whereas the patch 1 mutant alone rescued the formation of very large LDs and showed intermediate growth on terbinafine plates (Fig. 3g-i; S6c).

**Figure 3:**
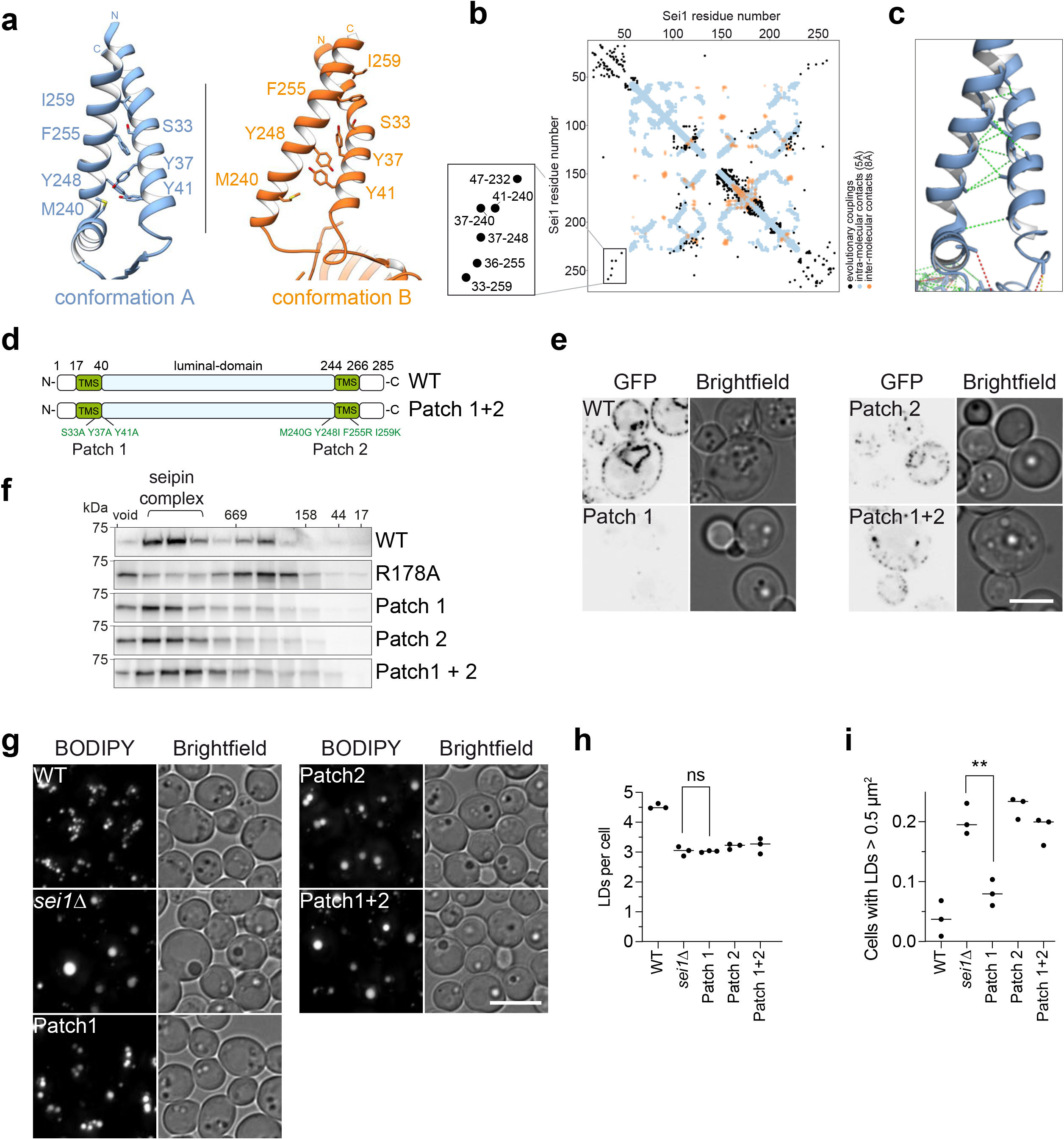
Transmembrane domain intramolecular interactions are important for seipin function and oligomer formation. **(a)** Detailed view of TM segments and switch regions in conformation A (blue) and B (orange), with residues indicating predicted intramolecular contacts. **(b)** Evolutionary coupling residues in yeast seipin highlight potential interactions in the TM segment regions. At the left, the membrane-embedded region is magnified. **(c)** Extended seipin structural model of conformation A, showing amino acids at least 10 residues apart in the primary sequence predicted to have beta-carbons interacting within 10Å distance, with maximal probability and over 70% probability mass, mapped onto the final model. Green dotted lines indicates that the actual distance is within 10Å, yellow within 12Å, and red for >12Å. View similar to Fig. 3a left side. **(d)** Overview of mutant constructs used in this figure. TMS, transmembrane segments. **(e)** Seipin TMD mutants integrate normally into the membrane and form WT-like foci. Seipin WT and indicated mutants expressed as C-terminal GFP fusion constructs from plasmids in *sei1Δ* cells. **(f)** Seipin intramolecular TMD mutants form normal oligomers. Size-exclusion chromatography of membrane extract in Triton X-100 from cells expressing *PGK1* promoter driven seipin and indicated mutants from the endogenous locus with C-terminal 13xmyc tag. Immunoblot with anti-myc antibodies. **(g)** LD morphology phenotype of strains expressing patch mutants from *PGK1* promoter. Densely grown cells were stained with BODIPY to visualize LDs. Size bar, 5 μm. **(h,i)** Quantification of experiment shown in g. One dot equals one separate experiment. **, p<0.01; ns, not significant.

Together with previous findings for human seipin^19^, our results highlight the importance of the seipin TM segments for LDAC function. Previously, it was reported that the yeast seipin TM helices are required for interaction with Ldb16^22^. Western blot analyses of cell lysates expressing the TM helix patch mutants or mutants with exchanged TM segments to FIT2 helices (Fig. S1) under control of the *PGK1* promoter showed that Ldb16 levels decreased in each of the TM segment mutants to a level generally similar to that in *sei1Δ* cells (Fig. S6d;^22^). However, much of Ldb16 expressed in the TM segment mutants appeared to be able to interact with seipin in pull-down assays (Fig. S6e), suggesting that seipin TM segments stabilize Ldb16 but are not strictly necessary for the interaction between the proteins.

Evolutionary coupling predicts similar intramolecular interactions between TM segments of fly and human seipin (S7a,b). To test whether the seipin TM helix architecture that we observed in our structure is conserved in evolution, we generated a series of chimeric proteins that contained portions of yeast seipin with regions of either fly or human seipin. Each of the mutants tested rescued yeast seipin deficiency to a similar extent as human or fly seipin (Fig. S7c), consistent with previous reports for human seipin ^22,29^. Furthermore, structural predictions of different seipin variants from different species by AlphaFold^30^ shows an architecture of the TM segments similar to the structure we resolved for conformation A (Fig.S7d). In summary, this suggests that the TM architecture is both critical for function and conserved through evolution.

### The Seipin Switch Region Is Required for Maintaining Seipin Complexes and Function

The main feature of the two TM segment conformations of the alternating subunits is that the TM helices of conformation B are tilted to the center of the seipin cage and interact with the neighboring TM helices of conformation A (Fig. 1c-e). This architecture is enabled by the flexibility of the switch regions that change most dramatically between conformation A and B. In particular, the switch region connecting to the seipin C-terminal TM segment showed a marked difference between the A and B conformations; it formed a kink in the A conformation but extended into a continuous α-helix with the C-terminal TM segment in conformation B (Fig. 1d; 4a; Suppl. Movie 1). This region also contained a highly conserved F_232_xxGLR sequence motif (Fig. S1a; S8a).

**Figure 4:**
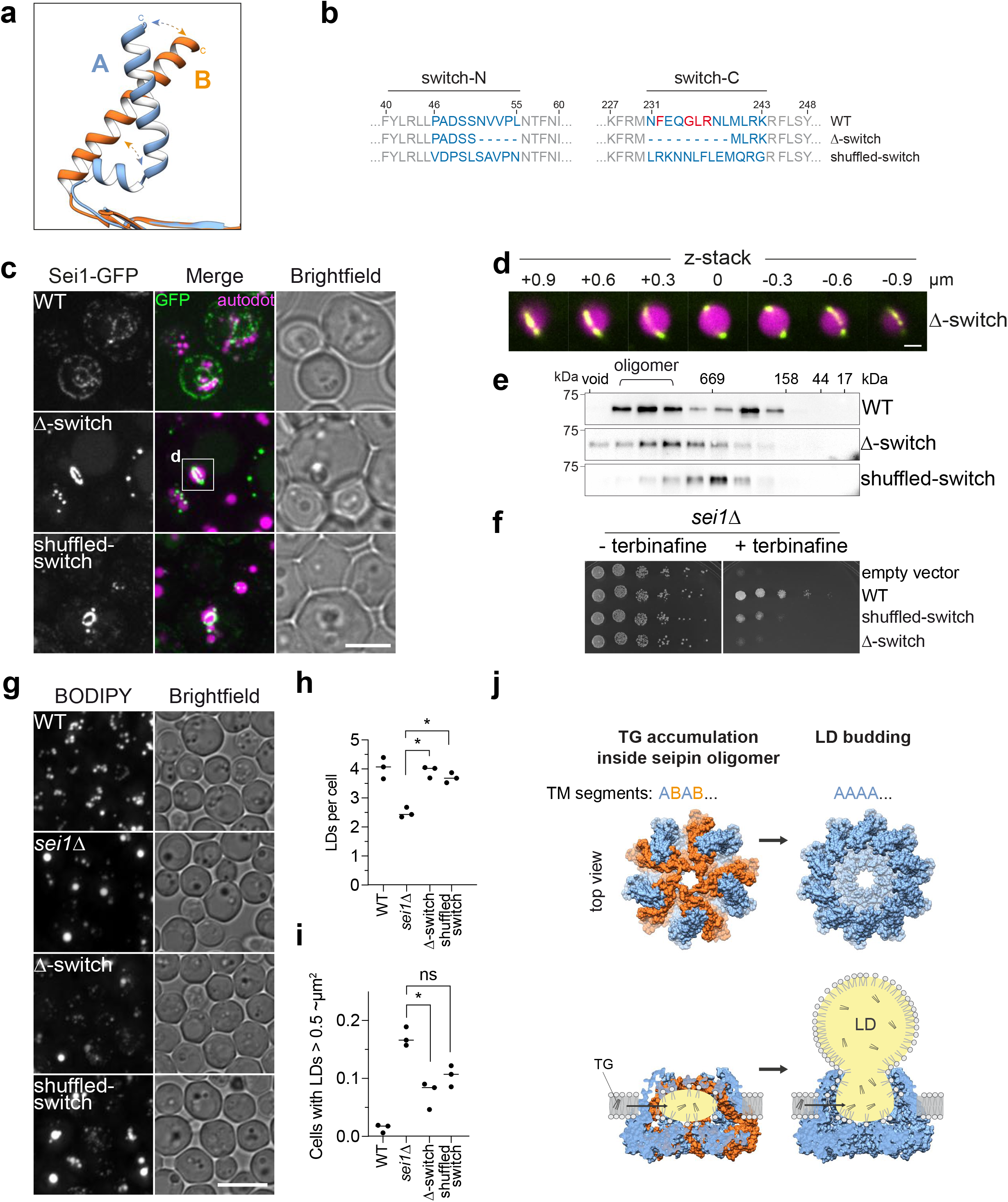
The seipin switch regions are required for seipin complex formation and function. **(a)** Detailed view of conformational change in C-terminal membrane helix comparing superimposed conformations A and B. Conformation A shows kinked alpha helix, and conformation B has an extended helix. **(b)** Overview of switch mutant constructs. **(c)** Seipin switch mutants forms large ring structures around LDs. Cells expressing C-terminal GFP-tagged seipin and indicated mutants from plasmids in *sei1Δ* cells. LDs were stained with autodot dye. Size bar = 5 μm. White box indicated area is shown in d. **(d)** Enlarged view and z-stack of seipin ring structures shown in Δ-switch mutant in c. Size bar, 1 μm. **(e)** Shuffled-switch mutant is unable to form WT-like oligomers in detergent extracts. Size-exclusion analysis of membrane extract from cells expressing *SEI1*-13xmyc or indicated mutants from the endogenous locus driven by integrated *PGK1* promoter. Immunoblot is shown. **(f)** Growth of yeast strain *sei1Δ* carrying plasmids with C-terminally GFP-tagged *SEI1* (WT), or indicated mutants on synthetic medium +/− 100 μg/ml terbinafine. **(g)** LD morphology analysis of strains shown in e. Size bar = 5 μm. **(h,i)** Quantification of LD morphology analysis shown in f. *, p<0.05; ns, not significant. **(j)** Model of seipin function in TG phase separation and LD budding by changing conformations of the transmembrane segments. Left side shows the conformation we obtained experimentally, and right side a predicted version of an “open” conformation based on all TM segments in the A conformation. Bottom model shows side views with TG accumulation in the complex.

To determine if the switch regions are important for seipin function, we deleted or shuffled their amino acid sequence (residues 46–55 and 231–244; Fig. 4b). The resulting shuffled-switch and Δ-switch mutants showed expression comparable to WT when expressed from the *PGK1* promoter (Fig. S8b). Disrupting the switch regions dramatically affected the cellular localization of the resulting protein, compared with WT. Instead of seipin foci commonly found at the contact site between the ER and LDs^13,22^, both mutants formed large rings that appeared to encircle large LDs, reminiscent of Saturn’s rings (Fig. 4c,d). The unusual pattern of switch mutant protein localization prompted us to hypothesize that these mutations weaken the interactions between TM segments of neighboring monomers by changing the arrangements of seipin’s A and B conformations. To investigate this possibility, we tested the prediction that complexes of seipin with shuffled switch regions are less stable in cells. We found that shuffled and Δ-switch formed smaller oligomers in detergent-solubilized protein extracts as analyzed by size-exclusion chromatography (Fig. 4e).

To test whether the switch regions of seipin are important for function, we assayed the ability of shuffled- and Δ-switch mutants to provide seipin function *in vivo.* Expression of mutant seipin versions with altered switch regions were unable to complement *sei1*Δ growth on terbinafine and only partially rescued the LD phenotype of *sei1Δ* cells (Fig. 4f-i).

## DISCUSSION

Understanding the function of seipin is crucial to deciphering the mechanism of LD formation from LDACs in the ER. Here we report a structural model for nearly all of the seipin protein of *S. cerevisiae* that combines a high confidence 3.2-Å molecular model based on cryo-EM of seipin’s lumenal domains, the switch regions, and TM segments, with an extended molecular model of the TM segments generated by an AI structure-prediction approach.

Core elements of the seipin structure appear to be evolutionarily conserved in yeast, fly, and human proteins^15,16,31^. The lumenal α/β-sandwich fold domain is well-resolved and has similar features in all species analyzed, except for the centrally located hydrophobic helix. Human and fly seipin have hydrophobic helices protruding into the center of the lumenal ring oligomer, whereas the analogous region in yeast consists of two short helices that are more hydrophilic. In human and fly seipin, the hydrophobic helix region is needed for interaction with LDAF1^19^ and has been proposed to interact with TG^17,18^. In yeast, however, we found that mutations of this region had little effect on seipin function. If an analogous central hydrophobic helix is also required in yeast, the yeast-specific Ldb16 protein could provide this function in trans for the LDAC. Because we found no density of Ldb16 in our yeast structure, our study cannot address this question.

The function of the lumenal domain remains uncertain. While this region was reported to bind anionic phospholipids^16^, whether this contributes to seipin function is unknown. Alternatively, the lumenal domain might primarily serve as a structural anchor for forming LDs, positioning key elements of the protein such as the hydrophobic helices at the membrane (for fly and human seipin) and the transmembrane helices at the budding neck. Although the yeast seipin complex contains 10 monomers, rather than 12 and 11 subunits in fly and human seipin, respectively^15,16^, the rings formed by the luminal domains of each species are similar in outer diameter and would provide similar diameters to necks of budding LDs.

An important feature of our yeast seipin model is the alternating conformations for monomeric subunit TM segments in the yeast decamer. The regions that change most between the two conformations are the switch regions, which are evolutionarily conserved between species (Fig. S1a; S8a). Consistent with our findings, *ab initio* structure prediction using the AI-system AlphaFold predicts that the TM segments of various metazoan seipins have a conformation similar to our experimentally determined structure of yeast seipin conformation A (Fig. S7d;^32^). Inasmuch as salient features of protein machines are most often conserved evolutionarily, we consider it likely that similar alternative conformations for the TM segments are possible for the human, worm, or fly proteins. However, although fly seipin with 12 monomers could adopt a symmetrical arrangement of A/B conformations, such symmetrical alternating conformations would be impossible for the 11-mer reported for human seipin^16^. This suggests that either human seipin complexes may be asymmetric, or that seipin can contain a mix of A and B conformations at any given time *in vivo*, such that symmetry in this respect is not important.

Considering our findings and data available from previous reports, we can propose a molecular model for seipin function during LD biogenesis. In this model, the seipin cage sits in the ER and excludes phospholipids from its central cavity to provide a space for TG molecules to interact with each other, rather than with phospholipid acyl chains. Neutral lipids such as TG, but not phospholipids with hydrophilic headgroups may diffuse into the complex through gaps in the plane of the membrane between seipin monomers. The net result is that seipin would allow interactions of TG molecules, thus catalyzing lens formation and growth. As the TG lens grows, the seipin oligomer may open towards the cytoplasm, with all subunits in the A conformation, and thus release the lens to generating an LD bud (Fig. 4j). As the forming LD grows, the TM segments could further tilt away from the center of the dome to accommodate the growing LDs. In agreement with this interpretation, our cryo-EM data suggest a higher degree of flexibility in the TM segments towards the cytoplasmic side of the seipin complex.

To maintain the neck of ER with LDs, and to allow this change in architecture, the switch region and interactions of TM segments would be particularly important. Consistent with this model, mutants in the switch region appear to lead to a seipin complex that cannot maintain a constricted neck at the ER-LD junction but rather dissociates and integrates more seipin subunits, eventually forming the large-diameter ring structures that we found around large LDs (Fig. 4c,d). Possibly related to such interactions, larger diameter rings of seipin form around LDs in *C. elegans*^33^.

Our model provides a conceptual framework for seipin function that can now be further tested by experiments and molecular modeling. It will be important to also integrate the structures and functions of additional known LDAC components, such as Ldb16 and the Ldo proteins in yeast, or LDAF1 in humans. Testing and refinement of the model should result in an increasingly clear understanding of this elegant protein machinery that governs the process of making oil droplets in cells.

**Table 1.** Cryo-EM data collection, refinement, and validation statistics.

|  | Sei1 in Pmal-C8 |
| --- | --- |
| **Data collection and processing** |  |
| Magnification | 105,000 |
| Voltage (kV) | 300 |
| Electron exposure (e-/Å<sup>2</sup>) | 28.73 |
| Defocus range | -1.2, -2.5 |
| Pixel size (Å) | 0.825 |
| Symmetry imposed | C5 |
| Map resolution (Å) | 3.2Å |
| FSC threshold | 0.143 |
| B factors (Å<sup>2</sup>) | -75 |
| **Refinement** |  |
| Initial model used (PDB code) | NA |
| Map sharpening B factor (Å<sup>2</sup>) | -75 |
| **Model composition (dimer model)** |  |
| Non-hydrogen atoms | 3830 |
| Protein residues | 470 |
| Water | 0 |
| Ligands | 0 |
| **Geometric deviations (RMSD)** |  |
| Bond length (Å) | 0.006 |
| Bond angles (°) | 0.742 |
| **Validation (dimer model)** |  |
| MolProbity score | 1.92 |
| Clashscore | 11.24 |
| Poor rotamers (%) | 0.23 |
| Ramachandran favored (%) | 94.81 |
| outliers (%) | 0 |
| Rama distribution Z-score | -1.77 +/- 0.39 |

## ACKNOWLEDGMENTS

We thank Siyoung Kim and Dr. Greg Voth (Department of Chemistry, University of Chicago) for critical discussions, Drs. Kelly Brock and Deborah Marks (HMS, Systems Biology) for advice on evolutionary coupling analyses, and Dr. Chao-Wen Wang (Institute of Plant and Microbial Biology, Academia Sinica, Taipei City) for kindly sharing Ldb16 antibody. We thank Dr. Danesh Moazed (HMS, Cell Biology) for sharing of equipment, Drs. Sarah Sterling, Richard Walsh and Zongli Li at the Harvard cryo-EM center for EM data collection, Joshua Reus, Nathan Paul, Adeline Gan, and Danny Chukwuma for generating and phenotyping several yeast strains, and Gary Howard for editorial assistance. This work was supported by NIH R01GM124348 (to R.V.F.), NIH R01GM084210 (to J.M.G.), a German Research Foundation (DFG) fellowship AR1164/1-1 (to H.A.), and an American Heart Association postdoctoral fellowship 18POST34030308 (to X.S.). T.C.W. is an investigator of the Howard Hughes Medical Institute.

## AUTHOR CONTRIBUTIONS

H.A., F.A.H., M.L., J.M.G., R.V.F., and T.C.W. designed and supervised the project. H.A., X.S., B.F., X.C., R.R., and J.M.G. carried out and analyzed experiments. H.A., and X.S. performed cryo-EM analysis and model building. C.A. and F.D. carried out machine learning calculations and modelling on the transmembrane helices. H.A. created figures and H.A., J.G., R.V.F., and T.C.W. wrote the manuscript, which all authors read and edited.

## COMPETING INTEREST STATEMENT

The authors declare no competing interests.

## SUPPLEMENTARY FIGURE LEGENDS

**Figure S1:**
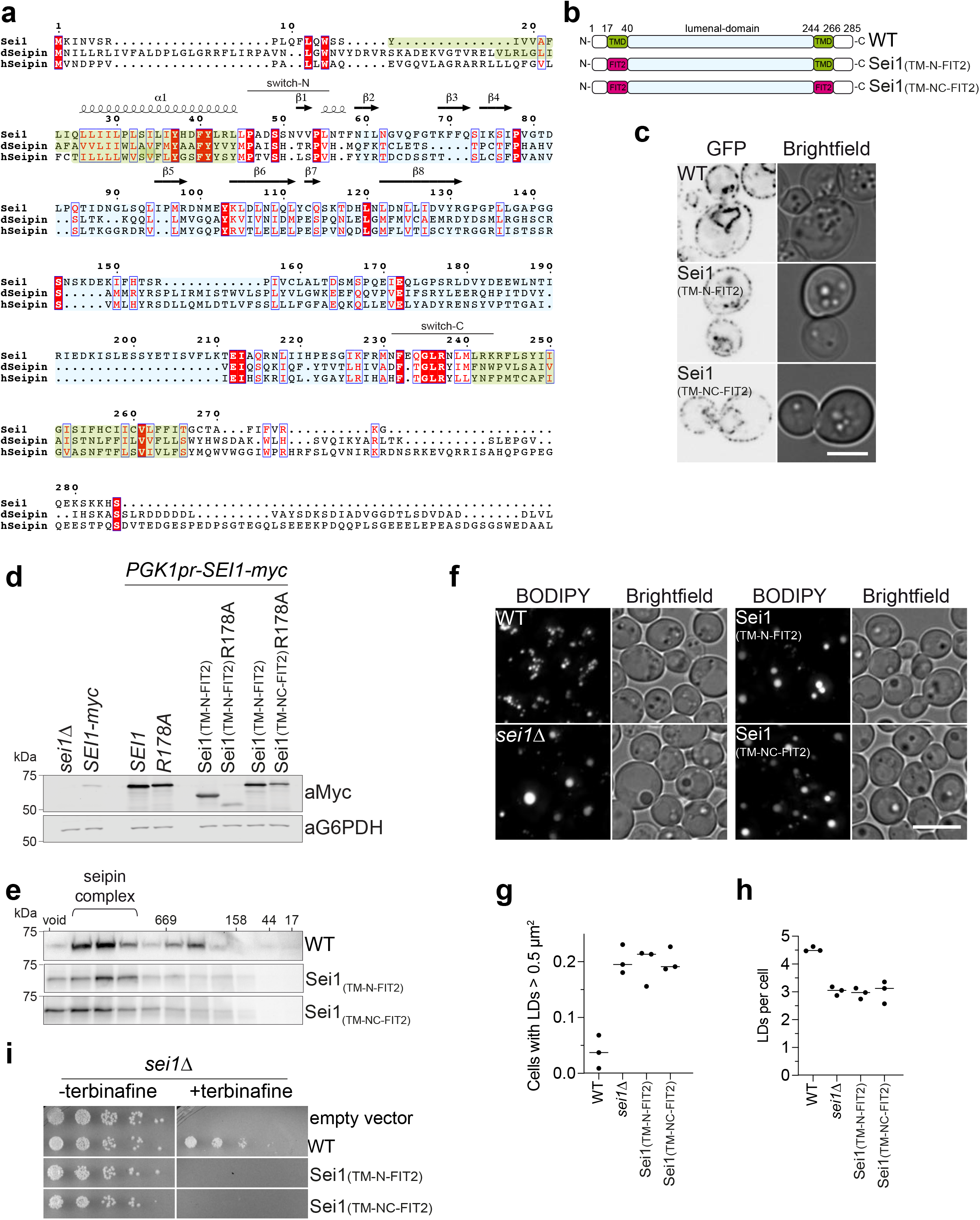
Transmembrane segments of seipin are conserved and required for function. **(a)** Sequence alignment of yeast (*Saccharomyces cerevisiae*) seipin (Sei1) protein sequence with *Drosophila melanogaster* (dseipin) and human seipin (hseipin) in T-COFFEE^34^, plotted in ESPript 3.0^35^. Identical residues are colored in red boxes, red characters and blue framed residues indicate similarity in a group or across groups, respectively. TM segments are colored in green and lumenal domains in cyan background similar to overview in b. **(b)** Overview of mutants analyzed in this figure. TMS, transmembrane segment. **(c)** Localization of seipin WT and mutant constructs expressed from plasmids in *sei1Δ* cells. **(d)** Expression level of WT and mutant constructs tagged with C-terminal 13xmyc. *SEI1*-myc indicates expression level from endogenous promoter. **(e)** Transmembrane mutants form normal oligomers in detergent extracts. Size-exclusion analysis of membrane extract from cells expressing *SEI1*-13xmyc or indicated mutants from the endogenous locus driven by integrated *PGK1* promoter. **(f)** Analysis of LD morphology using BODIPY staining. Seipin mutants with C-terminal 13xmyc tag were expressed from *PGK1* promoter. Size bar, 5 μm. **(g,h)** Quantification of experiment in panel f. **(i)** Growth of indicated mutants on synthetic medium +/− 100 μg/ml terbinafine.

**Figure S2:**
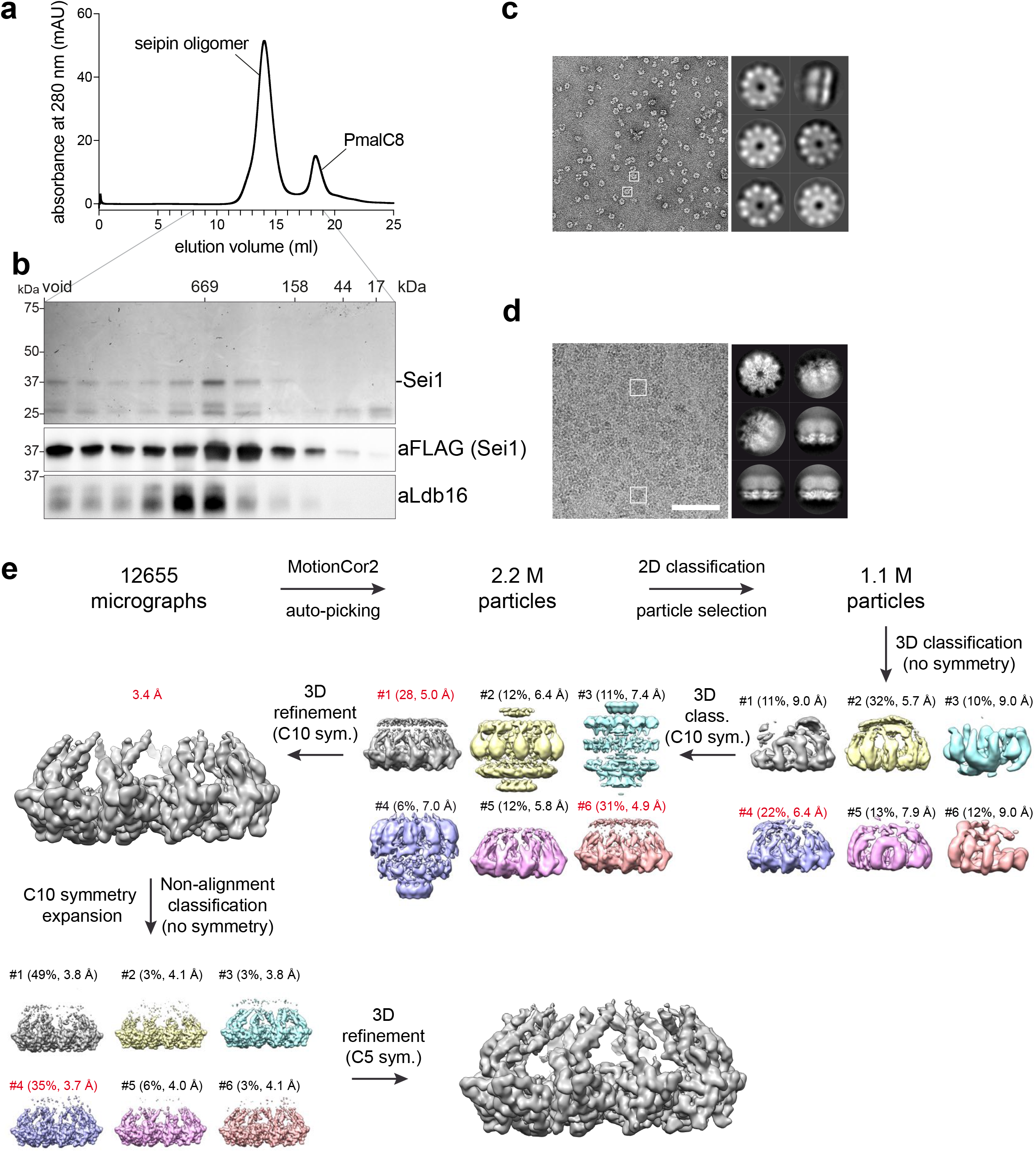
Purification and cryo-EM image processing of yeast seipin-Ldb16 complex. **(a)** Sei1-Ldb16 complex was purified from yeast as described in Experimental Procedures. After extraction of the complex in Triton X-100, detergent was subsequently exchanged to digitonin, and finally to PmalC8. The complex was separated by size-exclusion chromatography column in buffer without detergents. **(b)** Analysis of 1-ml fractions (8-18 ml) after SDS-PAGE by Coomassie Blue staining (top) or Western-blot (bottom). **(c)** Representative negative stain-EM image of purified complexes shown in a and b. Right side shows 2D class averages. White boxes indicate single oligomers. **(d)** Representative cryo-EM image of purified Sei1-Ldb16 complex. Right side shows 2D class averages. White boxes indicate single oligomers. Size-bar, 500 Å. **(e)** Three-dimensional classification and refinement of cryo-EM particles in Relion 3.0.

**Figure S3:**
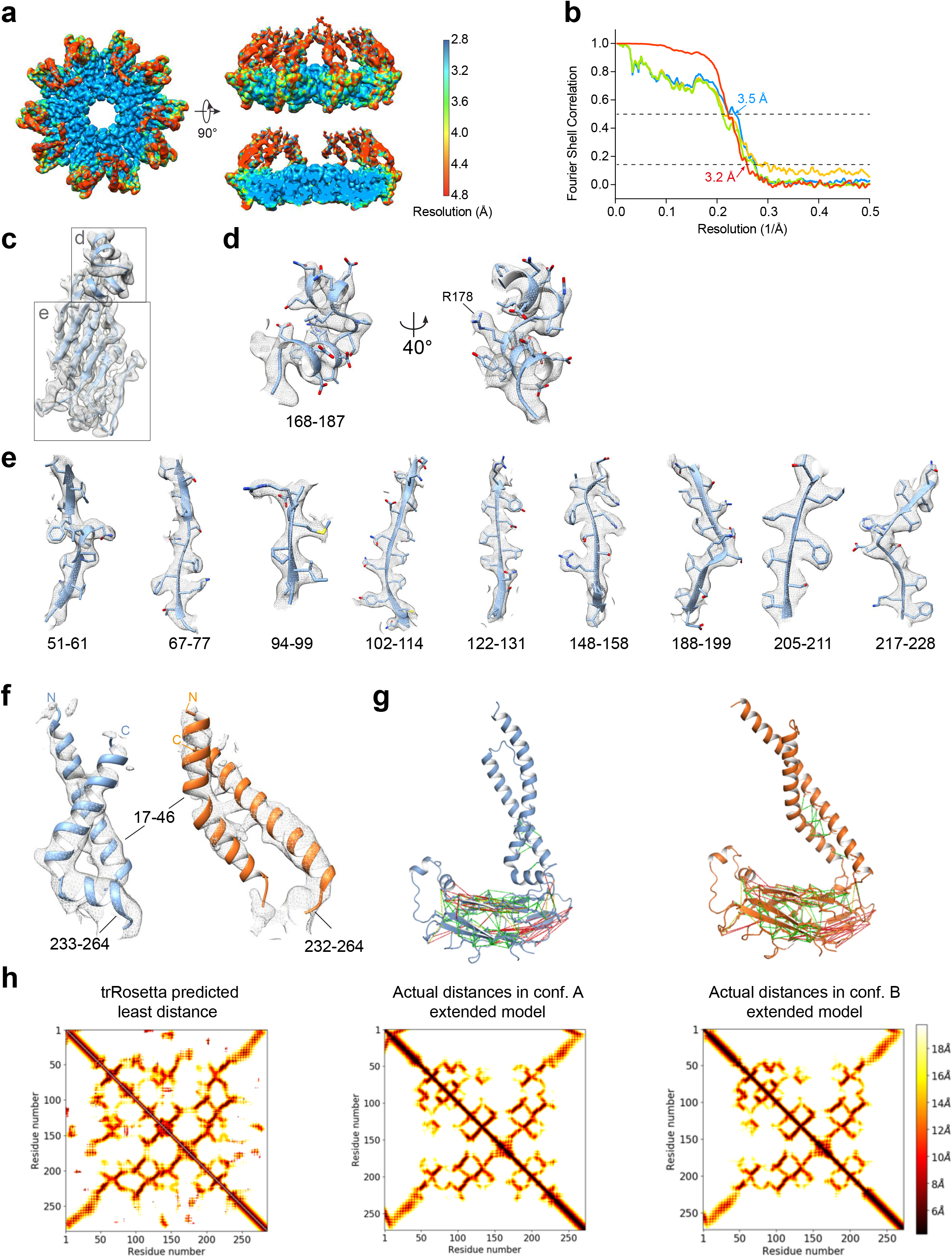
Single-particle cryo-EM analysis of Sei1-Ldb16 complex. **(a)** Local resolution mapped onto EM density map using Resmap^36^ shows differences between lumenal and transmembrane regions of the map. **(b)** FSC curves: gold-standard FSC curve between the two half maps with indicated resolution at FSC = 0.143 (red); half-map 1 (green), half-map 2 (orange) and the atomic model refined against half map 1 (blue). **(c-e)** Superimposed cryo-EM densities from sharpened map with atomic model for central alpha-helices (d) and individual beta-sheets (e). **(f)** Superimposed cryo-EM densities from unsharpened map with atomic model for TM segments of conformation A (blue) and conformation B (orange). **(g)** Extended models for conformation A (left) and B (right). Residues at least 10 residues apart in the primary sequence predicted to have beta-carbons interacting within 10 Å distance, with maximal probability and over 70% probability mass, mapped onto the final model of conformation A (left) and B (right). Green indicates that the actual distance is within 10Å, yellow within 12Å, and red for >12Å. **(h)** The predicted and actual distances between β-carbons of residues in the seipin monomer. The color of each pixel corresponds to the distance in Å between these atoms. Plotted on the left is the least distance predicted by trRosetta for each pair of CB atoms. In the middle are actual distances in conformations A, and conformation B (right). The trRosetta pipeline correctly predicts interactions between the N- and C-terminal helices for both conformations (from residues 10–40 and 250–280).

**Figure S4:**
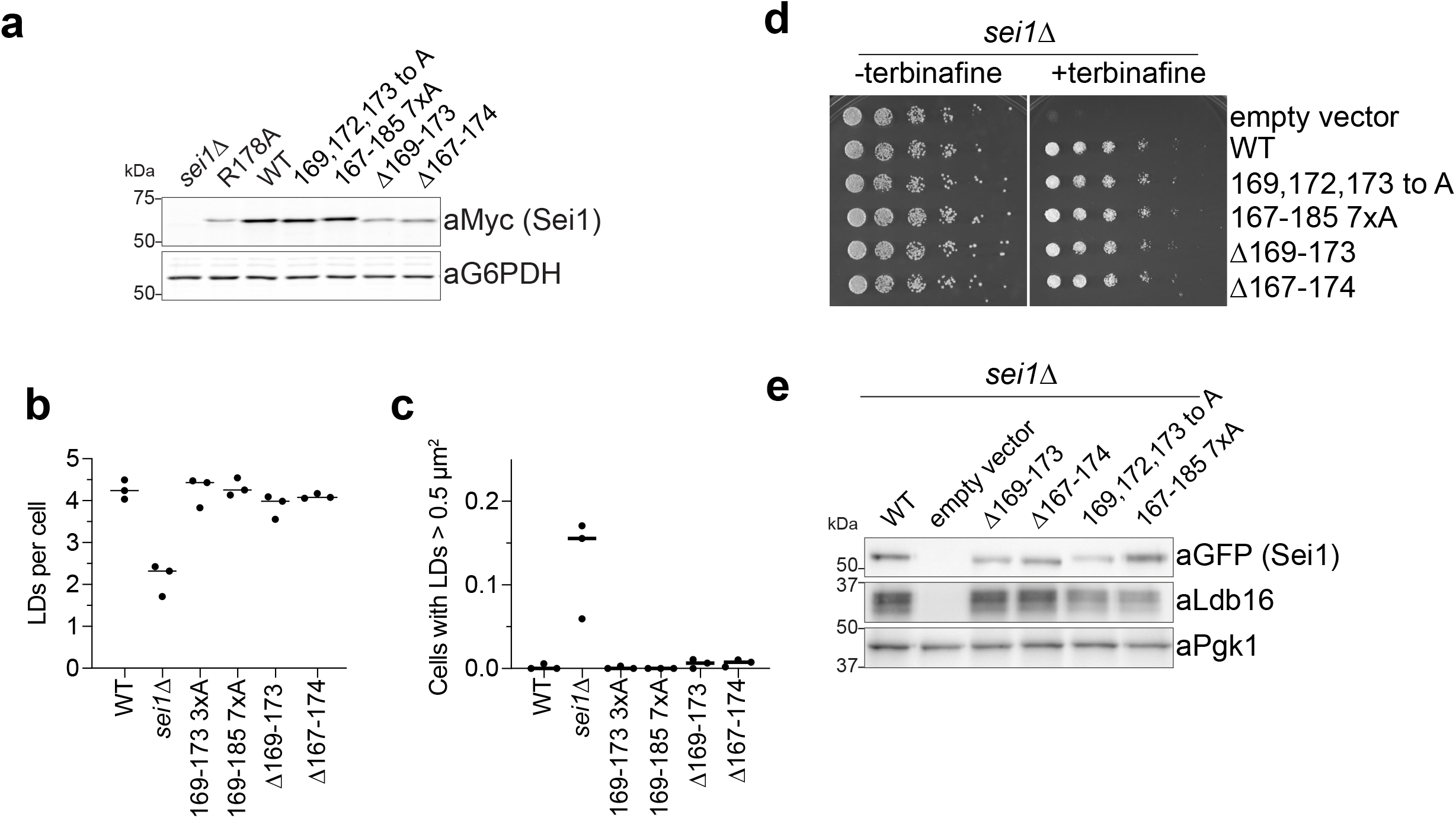
Mutants in seipin’s lumenal central helix retain function in vivo. **(a)** Western blot analysis of seipin expression level. Cells expressing WT or indicated mutant constructs with C-terminal 13xmyc tag from the endogenous promoter. Sei1 detected with anti-myc antibodies. **(b,c)** Quantification of images shown in Fig. 2c. **(d)** Growth of yeast strain *sei1Δ* carrying vectors with C-terminally GFP-tagged *SEI1* sequences or empty vector on synthetic medium +/− 100 μg/ml terbinafine. **(e)** Western blot analysis of whole-cell-lysates from strains in d using antibodies against GFP to detect seipin, against Ldb16 or Pgk1 as loading control.

**Figure S5:**
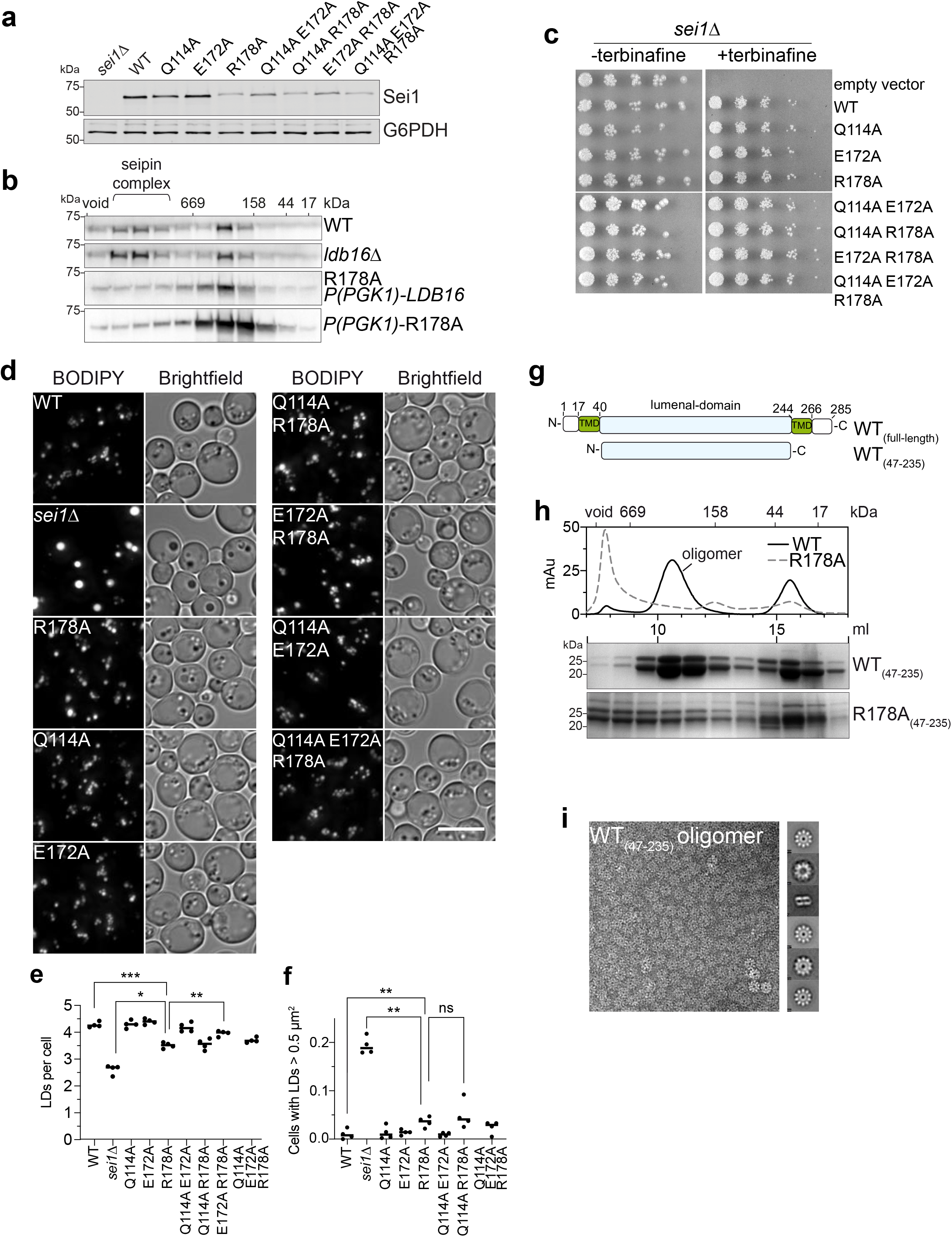
Lumenal domain interactions are mediated by R178. **(a)** Western blot analysis with anti-myc antibodies of lysate from strains expressing WT seipin or indicated point mutations from the endogenous locus with C-terminal 13xmyc tag. **(b)** Size-exclusion chromatography of Triton X-100 solubilized membrane extracts of indicated strains expressing C-terminal 13xmyc-tagged seipin. **(c)** Growth of yeast strain *sei1Δ* carrying vectors with C-terminally GFP-tagged *SEI1* mutants or empty vector on synthetic medium +/− 100 μg/ml terbinafine. **(d)** LD morphology of strains expressing indicated seipin mutants with C-terminal 13xmyc from endogenous locus. Size bar, 5 μm, **(e,f)** Quantification of experiment shown in d. *, p<0.05; **, p<0.01; ***, p<0.001, ns, not significant. **(g)** Overview of lumenal domain construct purified from *E. coli*. **(h)** Size-exclusion chromatography analysis of affinity purified WT lumenal domain (WT(47-235)) or R178A(47-235). Top, traces of absorbance at 280 nm in mAu of WT and R178A lumenal domains. Bottom, SDS-PAGE analysis of 1-ml fractions by Coomassie staining. **(i)** Negative stain-EM analysis of WT lumenal domain oligomers shown in h. Right side shows 2D class averages.

**Figure S6:**
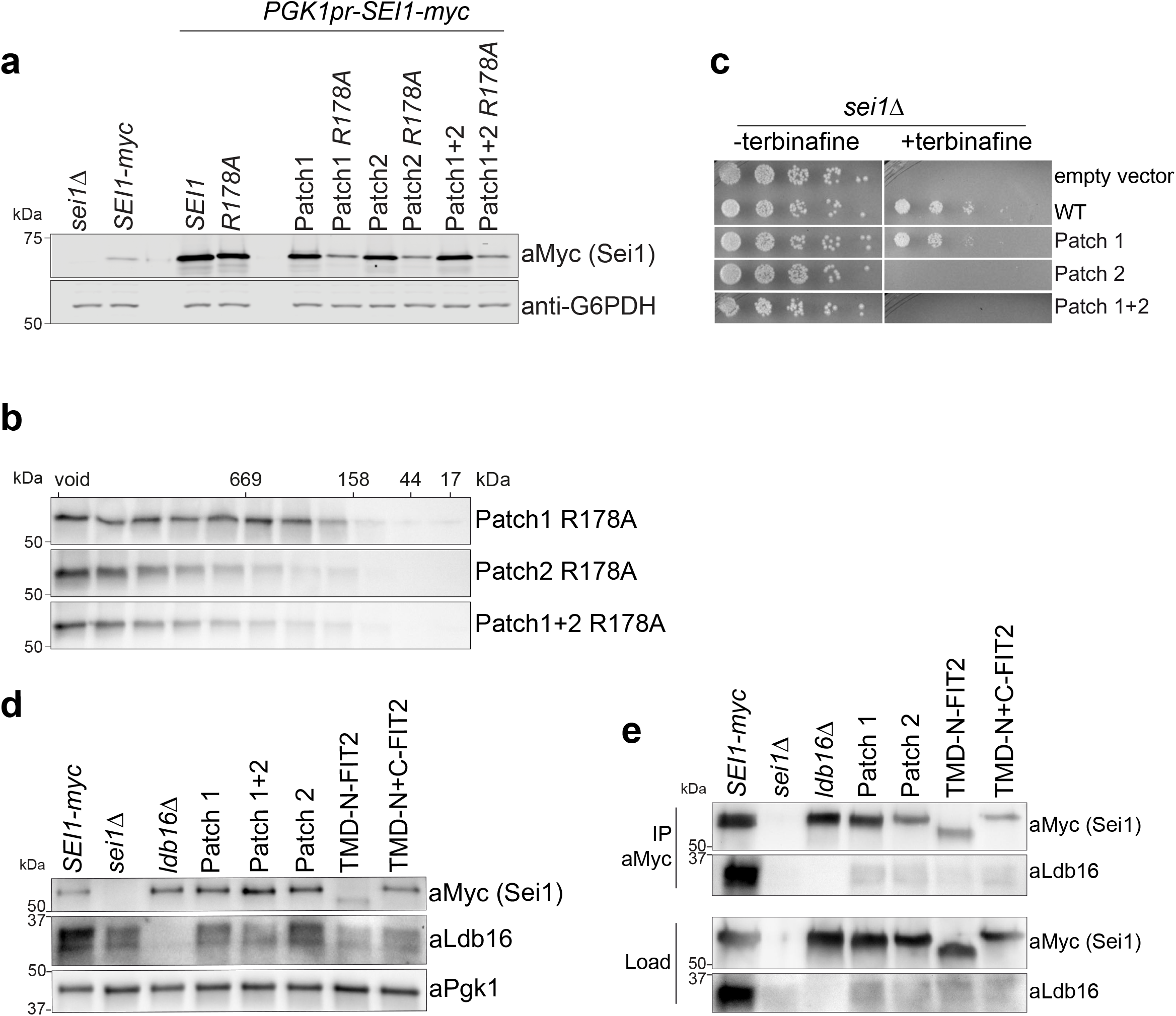
Intramolecular transmembrane segment interactions are crucial for seipin function. **(a)** Western blot analysis of whole-cell lysates from strains expressing C-terminal 13xmyc tagged seipin variants from endogenous or *PGK1* promoter. **(b)** Western blot analysis of fractions from size-exclusion chromatography of Triton X-100 solubilized membrane extracts carrying indicated mutations with C-terminal 13xmyc tag. **(c)** Growth of yeast strain *sei1Δ* carrying plasmids with C-terminally GFP-tagged *SEI1* from yeast (WT), or indicated mutants on synthetic medium +/− 100 μg/ml terbinafine. **(d)** Western blot analysis of whole-cell lysates from strains expressing indicated seipin mutants under control of the *PGK1* promoter and C-terminal 13xmyc tag. **(e)** Immuno-precipitation of indicated seipin mutants via anti-myc resin. Equal amounts of load (detergent solubilized membranes in Tx100) and eluate fractions were loaded.

**Figure S7:**
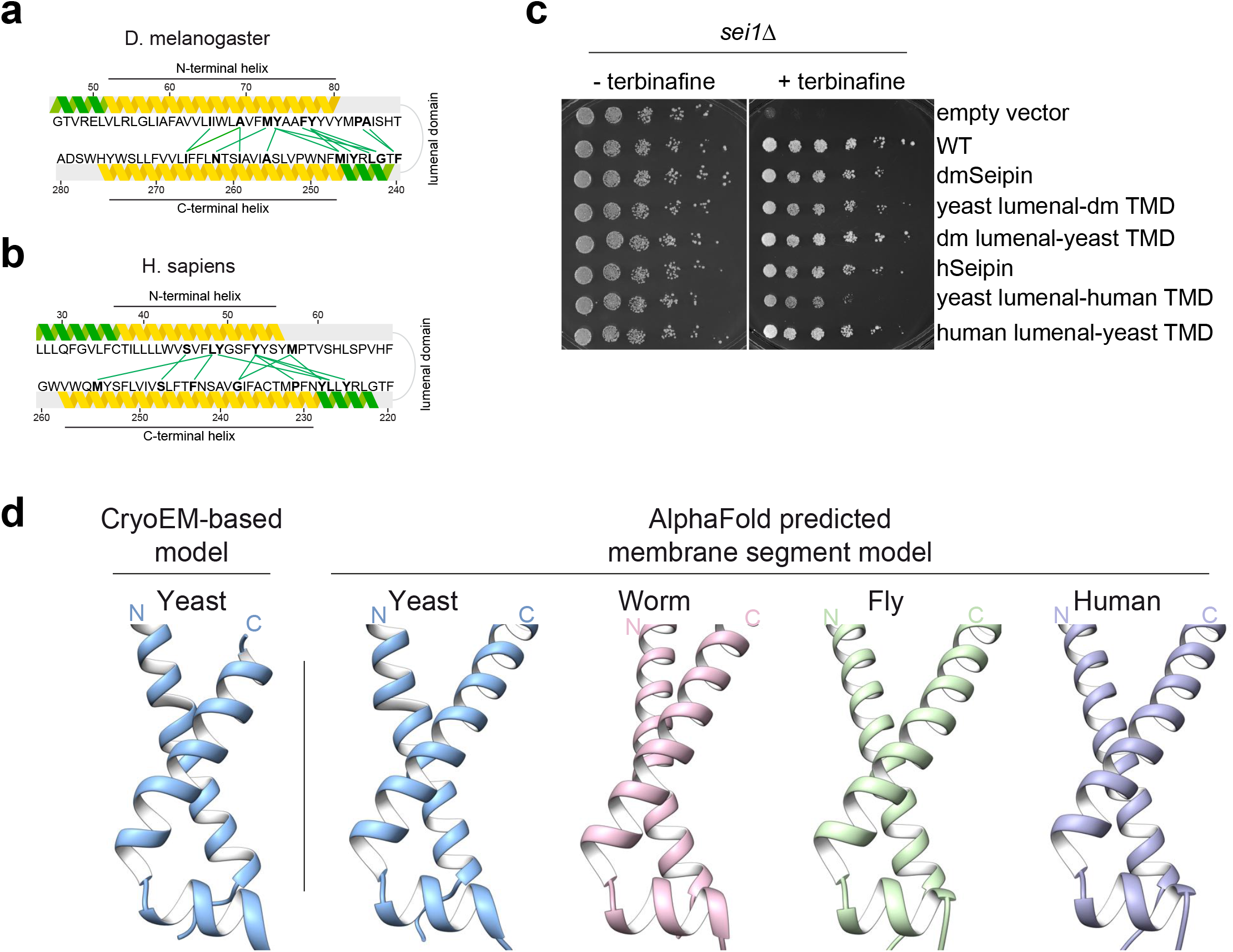
Transmembrane segment architecture is conserved. **(a,b)** Highest ranking evolutionary couplings (green lines) within seipin transmembrane and switch regions mapped onto (a) *D. melanogaster* or (b) human sequences. Yellow and green helices indicate secondary structure prediction by Phyre2^37^ of membrane embedded or hydrophilic helices, respectively. Coupling residues are indicated in bold. **(c)** Growth of yeast strain *sei1Δ* carrying plasmids with C-terminally GFP-tagged *SEI1* from yeast (WT), *D. melanogaster* (dmSeipin), human (hSeipin) or chimeric constructs on synthetic medium +/− 100 μg/ml terbinafine. **(d)** The architecture of seipin transmembrane helices is predicted to be conserved. Comparison of switch and transmembrane regions of our structural model (left) with predicted structure of yeast (*S. cerevisiae*); worm (*C. elegans*), fly (*D. melanogaster*) or human by AlphaFold^38^.

**Figure S8:**
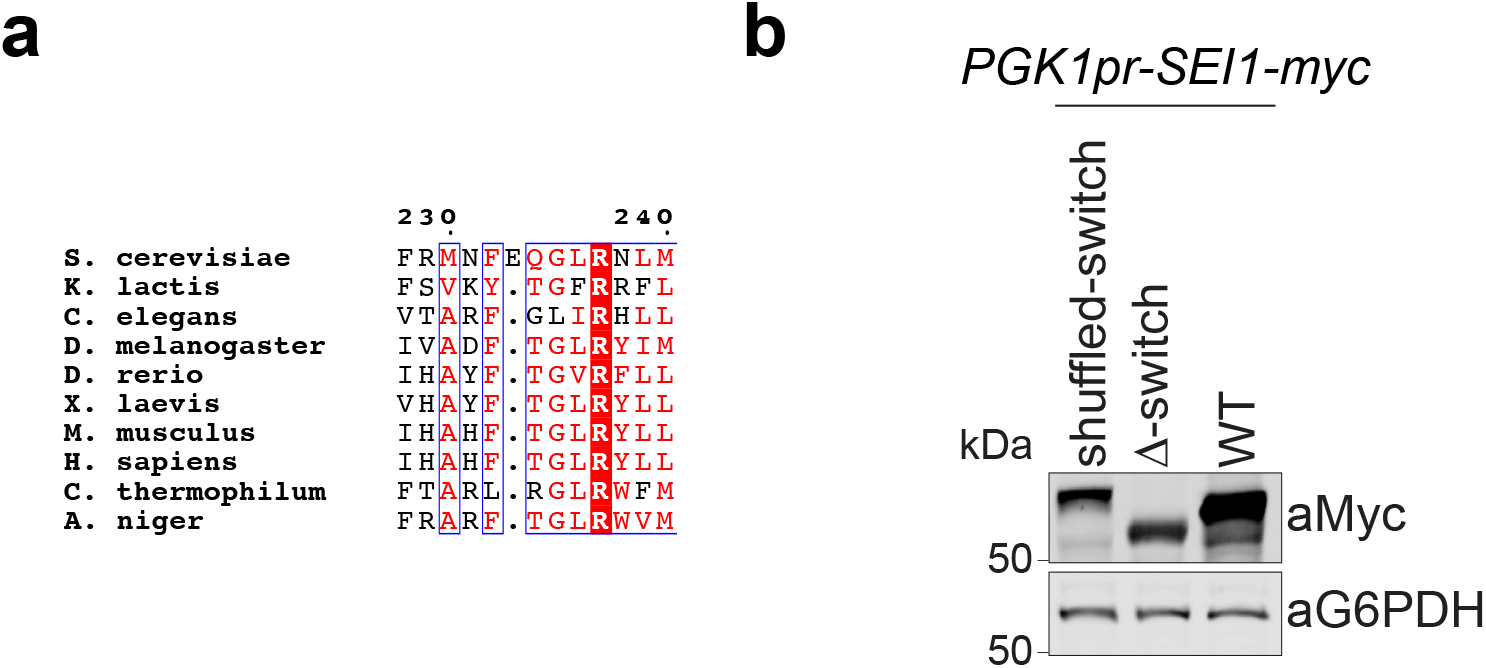
Switch regions are required for seipin function. **(a)** Sequence alignment of seipin sequences from different species shows conserved F_232_xxGLR sequence motif. Identical residues are colored in red boxes, red characters and blue framed residues indicate similarity in a group or across groups, respectively. **(b)** Western blot analysis of whole cell lysates from strains expressing indicated switch mutants or WT seipin under control of the *PGK1* promoter with C-terminal 13xmyc tag.

## EXPERIMENTAL PROCEDURES

### Plasmids

All plasmids used in this study are listed in Table S1. A yeast shuttle vector, pBMF1, a derivative of pRS313, was constructed that contained the following in tandem, flanked by HindIII and SacII sites: 207 bp *SEI1* 5′ untranslated region (UTR), *SEI1* coding region in frame with 13 copies of sequence encoding the myc epitope, *ADH1* terminator, the pRS313 *HIS3* cassette, and 236 bp of *SEI1* 3′ UTR. *SEI1* mutants were generated in pBMF1 using appropriate PCR products and NEBuilder HiFi DNA Assembly (New England Biolabs).

A plasmid, pSK^-^-NAT-PGK, was constructed containing the nourseothricin (NAT) resistance cassette (from pFA6-natMX6) and the *PGK1* promoter (982 bp of 5′ untranslated sequence). For overexpression of seipin mutants, a PCR product containing the NAT-PGK1 fragment was inserted upstream of the seipin coding region in the genome.

For expression of C-terminal GFP-tagged constructs, *SEI1* or gene-synthesized mutants (Integrated DNA Technologies) flanked by HindIII and BamHI were inserted into a pRS416 vector containing *ADH1* promoter (705 bp) and GFP. pHA234 expressing GFP alone served as empty vector control.

All PCR-derived fragments were fully sequenced in plasmids and mutations in the genome verified.

### Yeast strains

All strains (Table S2) were based on a W303-1A or BY4741 background. *PLN1* was knocked out when indicated by replacement with a hygromycin-resistance cassette. Seipin mutants were generated by transforming yeast with a the HindIII-SacII DNA fragment from pBMF1 (with appropriate mutations) containing *SEI1* sequences and the *HIS3* marker for selection of transformed clones. Homologous recombination was confirmed by PCR and mutations in the genome confirmed by sequencing.

Insertion of *GAL1* promoters and C-terminal 3xFLAG-TEV-2xProteinA tag to generate HAY60 was carried out by integration of PCR products from plasmids pYM-N22, pYM-N23^39^ and pFA6a-hphMX-(3×FLAG)-TEV-ProtA (gift from Michael Nick Boddy, Addgene plasmid # 52692).

For antibiotic selection, strains were selected on yeast peptone 2% dextrose (YPD) plates containing nourseothricin (GoldBio), hygromycin B (Thermo Fisher Scientific), or kanamycin (Sigma-Aldrich). Transformants were first grown overnight on YPD plates before stamping onto antibiotic plates or directly plated onto antibiotic plates after incubation in YPD shaking culture for 3 h at 30°C.

### Protein expression and purification

Sei1-Ldb16 complexes were expressed from yeast strain HAY60 grown in yeast peptone media supplemented with 2% galactose (YPG) for at least 24 h at 30°C in 1 L cultures. Densely grown cells were harvested by centrifugation, were washed 1x with water, and buffer A (50 mM Tris pH8.0, 150 mM NaCl, 0.5 mM EDTA, 10% glycerol). Cell pellets were resuspended in a small volume of buffer A supplemented with 35 μl/ml yeast Protease Inhibitor Cocktail (Sigma) and snap frozen in liquid nitrogen. Frozen cell pellets were lysed in a cryo-mill, and ground lysate powder was stored at −80 °C. For large purifications, typically 100 g of powder from ~10-L cultures was thawed at RT, supplemented with buffer A, and followed by centrifugation at 4000 g for 10 min to remove cell debris. Membranes were isolated by ultracentrifugation at 125,000 g for 1 h at 4°C, were resuspended in buffer A containing 1% Triton X-100 for 1-2 h at 4°C and centrifuged again for 1 h at 125,000 g. The supernatant was incubated for 2 h with 6 ml washed IgG Sepharose 6 Fast Flow beads (Cytiva) at 4°C on a nutator. Beads were washed with 10 ml buffer B (50mM Tris pH 8.0, 150 mM NaCl, 5 mM MgCl2,) + 0.05 % Triton X-100, 2x in same buffer + 0.5 mM ATP, 2x 6 ml of buffer B without detergent, and 6 ml of buffer B with 0.1% digitonin. Sei1-Ldb16 complexes were eluted by TEV-cleavage using home-made TEV protease in 3 ml of buffer B + 0.1% digitonin overnight at 8 °C with constant shaking (350 rpm). The eluate was concentrated in 100-kDa filters (Amicon) and separated on a Superose 6 Inc column in buffer B + 0.05% digitonin. Protein-containing fractions were combined and concentrated to 1.5-ml volume, and 1:3 (w/w) PmalC8 (Anatrace) was added. Mixture was loaded to 35 kDa dialysis filters in 50-ml falcons to buffer B, supplemented with 500 μl of Bio-Beads SM-2 (Bio-Rad) overnight at 4°C on a nutator. PmalC8-reconstituted protein complexes were subjected to another size-exclusion chromatography on Superose 6 Increase column in buffer B (Fig.S2a) and used for negative staining or cryo-EM sample preparation.

WT and R178A seipin lumenal domains were expressed in SHuffle T7 Express *E.coli* cells using plasmids pHA147 (WT_(47-235)_) and pHA144 (R178A_(47-235)_) that contained a C-terminal 6xHis tag. After induction at OD600 = 0.8 with 0.5 mM isopropylthio--β-galactoside and incubation at 16°C overnight, cells were harvested and lysed in buffer C (50 mM Tris pH8.0, 400 mM NaCl2, 5 mM MgCl2) supplemented with 1 mM phenylmethylsulfonyl fluoride and 20 mM imidazole in a Microfluidizer LM 20 (Microfluidics) run at 18,000 PSI. Lysate was cleared by centrifugation for 30 min at 20 000 g at 4°C, and supernatant was incubated with Ni-NTA agarose beads for 1 h at 4°C. Beads were collected and washed with buffer C + 5% glycerol and 40 mM imidazole, followed by elution in buffer C + 5% glycerol and 500 mM imidazole. Purified proteins were analyzed by size-exclusion chromatography using Superdex 200 Increase column in buffer C + 5% glycerol.

### Size-exclusion analysis of membrane extracts

Yeast strains expressing *SEI1*-13xmyc were grown in 25 ml of YPD culture at 30°C for 16–24 h were harvested and washed with water by centrifugation at 4000 g, 5 min. Cell pellets were resuspended in 600 μl of buffer A with protease inhibitor cocktail. 250 μl of 0.5-mm silica beads were added, and cells lysed in a bead beater 2x 30 s at full speed with 10 min breaks on ice. Lysate was harvested by centrifugation (20 s, 18,000 g) and precleared (5000 g, 10 min, 4°C). Membranes were collected by centrifugation at 125,000 g for 1h and solubilized in buffer A containing 1% Tx100 similar to sample preparation for Sei1-Ldb16 protein purification. Solubilized membranes (in typically 900 μl volume) were centrifuged again 1 h, 125 000 g, 4°C, and 500 μl were filtered in 0.2-μm filters and analyzed by size-exclusion chromatography on Superose 6 Inc column as described above, followed by SDS-PAGE, western blot and detection of myc tag using anti-myc monoclonal 9E10 (Thermo Fisher Scientific) and secondary antibodies anti-mouse-HRP (Cat# sc-516102; Santa Cruz Biotechnology).

### Immuno-precipitation

Solubilized membrane extracts in 1% Triton X-100 from 25 ml of YPD cultures were prepared as described above in 1.2 ml total volume and were added to 250 μl anti-myc agarose slurry (Thermo Fisher Scientific) in 2 ml tubes. After incubation for 1 h, 4°C on a nutator, beads were washed with 2x 1 ml buffer A + 0.01% Triton X-100. Bound proteins were eluted by addition of 50 μl Laemmli buffer and incubation for 30 min at 95 °C. Samples were analyzed by SDS-PAGE and western blot.

### EM sample preparation and data acquisition

Negative-stained samples were prepared as described40 and imaged on a Tecnai T12 microscope (Thermo Fisher Scientific) equipped with 4k x 4k CCD camera (UltraScan 4000; Gatan).

Cryo-EM samples were concentrated to ~3.5 mg/ml in 100-kDa filters, and 2.5 μl of sample was added to 30 s glow discharged Quantifoil holey carbon grids (Cu R1.2/1.3; 400 mesh), blotted with Whatman #1 filter paper with ~100% humidity and plunge frozen in liquid ethane using a Vitrobot Mark IV (Thermo Fisher Scientific). Images were collected on a Titan Krios electron microscope (Thermo Fisher Scientific), details see Table 3.

### EM data processing

Cryo-EM data processing was carried out as described previously^15^. Briefly, images were drift corrected by MotionCor2^41^ and binned 3 x 3 by Fourier cropping to a pixel size of 2.475 Å. Defocus values were determined using CTFFIND4^42^ and motion-corrected sums without dose-weighting. Motion-corrected sums with dose-weighting were used for all other steps of imaging processing. After particle picking, 2D classification of selected particles was performed in samclasscas.py. Initial 3D models of a cylindrical density matching the overall Fld1/Sei1 complex dimension were generated using SPIDER to perform the initial 3D classification. 3D classification and refinement were performed in Relion 3.0^43,44^ initially without application of symmetry. After the first rounds of 3D classification without symmetry on binned particles, the second round of classification was performed on selected particles without binning. This step was followed by global refinement on selected particles with C_10_ symmetry. Afterwards, the refined particle stack underwent symmetry expansion with C_10_, and was further classified without global angle search (non-alignment classification). In this and the following steps, the density model from previous refinement result was used as reference. For the final round of refinement C_5_ symmetry was imposed to generate the cryo-EM map of Sei1 showing signal of the transmembrane region. The final EM density map was sharpened by application of −75 B factor with the filtered resolution of 3.75Å by the program bfactor.exe^45^. Local resolution variation of EM density maps was calculated in ResMap 1.1.4^36^.

### Model building and refinement

Seipin density maps in MRC/CCP4 format were converted to structure factors MTZ format in PHENIX suite^46^. Models were built manually in COOT^47^ starting from the high-resolution region in the ER lumenal region, and iteratively refined in PHENIX real-space refinement procedure, followed by visual inspection and manual refinement in COOT. The transmembrane segments (residues 25-46; 234-258) of conformation A were manually built. Other parts of the TM segments of conformation A and B were modeled as described below.

### Molecular modeling of the transmembrane helices

The trRosetta^27^ neural network was run on the full-length sequence to generate 2003 aligned sequences. Filtering by 90% maximum pairwise sequence identity and 50% minimum sequence coverage yielded 921 sequences, which were used to derive pairwise constraints across the whole structure. The trRosetta constraints were input alongside density data to the Rosetta comparative modeling (RosettaCM)^48^. We leveraged the manually built model from residues 25-258 in the A conformation and 49–233 in the B conformation as starting models in this pipeline. 10,000 modelling trajectories were sampled for each conformer, and the top models selected by Rosetta showed good agreement to the density. These conformers were input as the asymmetric unit in Rosetta symmetric refinement. C_5_ symmetry was used to generate the final 10-mer model.

### Terbinafine growth assays

Yeast strain BY4741 *sei1Δ* was transformed with plasmids expressing seipin constructs from ADH1 promoters and C-terminal GFP tag by selection on synthetic medium without uracil. Cells were grown to early stationary phase for 16–24 h in 3-ml cultures in synthetic medium without uracil + 2% dextrose. OD600 was determined, and cells diluted to OD 0.25. Serial 1:5 dilutions in sterile water were performed in 96-well plates, and 3 μl were spotted onto plates with or without 100 μg/ml terbinafine (Sigma-Aldrich). Plates were imaged after 3–7 days of incubation at 30°C.

### Fluorescence microscopy

One μl BODIPY 493/503 from a 1 mg/ml stock in DMSO (stored dark) was added to 1 ml of culture in a 1.5 ml microfuge tube and incubated on a rocker (dark) for 30 min, centrifuged for 1 min at 2000 x g, and 950 μl of the supernatant removed. Cells were resuspended in the remaining media and 1.7 μl of the cell suspension applied to a slide for microscopy. Alternatively, cells from 3 ml culture were centrifuged (20 s, 18 000 g), resuspended in 50 μl synthetic medium + 5 μl of 1:250 diluted autodot dye (Abcepta), and incubated as described above.

The microscope hardware, and image acquisition and projections from z-stacks were as reported previously^49^, except the z-stack consisted of 25 images taken 0.35 microns apart, and Slidebook version 6.0.4 (Intelligent Imaging Innovations) was used. Alternatively, cells were imaged on a Nikon Eclipse Ti inverted microscope equipped with CSU-X1 spinning-disc confocal scan head (Yokogawa), 405-, 488-, and 561-nm laser lines, 100x Apochromat total internal reflection fluorescence 1.4 NA objective (Nikon), Zyla 4.2 Plus sCMOS, or iXon897 electron-multiplying charged-coupled device cameras (Andor) and NIS Elements AR software (Nikon).

### Cell culture

Typically, seipin protein expression was determined on cultures that were also subjected to fluorescence microscopy to determine number and size of LD. A colony from each strain was inoculated into 5 ml of SCD-defined medium^11^ and incubated for 20-24 h in a rotary shaker, then back-diluted to OD_600_ of 0.1/ml into 50 ml of SCD and incubated for 24 hours. The culture was then immediately processed for both fluorescence microscopy and immunoblotting.

### Cell segmentation

To facilitate cell segmentation in brightfield images, a deep learning pipeline for automatic instance segmentation was implemented, mostly following^50^. In short, we trained a convolutional neural network to jointly make three pixel-wise predictions: A seed map, a scalar bandwidth, and two-dimensional spatial embeddings, which were used to differentiate cells. After adding the pixel coordinates, the spatial embeddings should be constant over each cell, while different cells should have distinct embeddings.

To train the neural network, the above condition was encouraged in an indirect manner: For every cell, the average embedding vector was computed and a soft mask was grown, using a Gaussian kernel of the average predicted bandwidth over that cell in the embedding space. Using a loss for binary classification, these soft masks were driven to match the binary ground-truth masks of the cells. In this we deviated from^50^ and used the Dice-Loss algorithm^51^ which works well with the class imbalance between foreground and background.

In the inference procedure, an instance segmentation was inferred using an iterative algorithm^50^. The pixel with the highest score in the seed map was selected as the seed, and all pixels whose spatial embeddings are sufficiently close to the embedding of the seed pixel were clustered as a predicted instance. This process was repeated, conditioning the selection of the seed pixel to the not yet assigned regions, until all foreground pixels (i.e., pixels with a seed score over 0.5) were assigned to an instance.

As postprocessing, we filtered out predicted instances whose size falls below a threshold of 300 pixels as well as those that touch the image border. Finally, the convex hulls of the predicted segments were converted to a list of FIJI/ImageJ regions of interest, on which the downstream analysis was performed.

To generate the necessary ground-truth data, 18 images were annotated. For each living cell (i.e., cells without dense cytoplasm) in those images, an ellipse was drawn in FIJI/ImageJ, which was converted to a pixel-wise mask for that instance. These masks were then combined to generate the label images required to train the neural network. Of the 18 images, we used 14 for training and 4 for validation.

As the architecture of our model we choose a variant of U-Net^52^ with additional residual connections^53^ at every scale in both the encoding and decoding branches. Each convolution was followed by a batch normalization layer ^54^.

The network was trained on 1024×1024 pixel crops of the annotated images, which were randomly flipped and rotated to augment the training data and thereby combat overfitting. We used the Adam optimizer^55^ with a learning rate of 10−4. The model was trained on a single graphics card, while predictions were computed on the CPU to simplify deployment.

Finally, the calculated cells perimeters for each field, which were converted to ImageJ ROI files. Software was downloaded from Github and installed on Macintosh computers; it is available at https://github.com/hci-unihd/YeastCellSeg for public use.

### Fluorescence image quantification

Seipin loss-of-function results in fewer and larger “supersized” droplets (or aggregates of small droplets). The number of supersized droplets in seipin mutants are enhanced in *pln1Δ* strains^49^ and for this reason, most experiments with seipin mutants were performed in a *pln1Δ* background.

The heterogeneity of LD sizes, number, and tendency of LDs to cluster presented a challenge for automated LD counting. An ImageJ routine was written to count particles per cell (ROI) in each field iteratively at decreasing lower threshold (upper threshold was set at maximal) starting at 20,000 at 2000 increments and ending at 2000. (At 20,000 threshold only the largest droplets were counted, whereas at 2000, very dim droplets were counted while the point-spread function of larger ones in clusters merged.) The droplet number for each cell was the maximal particle count over the threshold range. This correlated well to droplet counts determined visually except the dynamic range was somewhat attenuated, as very dim droplets were counted (increasing the count), but LDs in clusters of droplets were not resolved (decreasing the count). However, the relative values among strains corresponded well to the visual counts (not shown).

Scoring cells with supersized droplets was performed by counting BODIPY-stained particles per cell at 10,000 lower threshold in ImageJ with an area (point-spread function at this threshold) of greater than 0.5 sq microns.

### Statistical analysis

For analysis of LD phenotypes, at least three independent experiments were performed with all mutant sets. Three microscope fields for each strain in each experiment, each typically with 200 cells, were analyzed as described for droplets per cell and cells containing LDs over 0.5 μm^2^. The mean value was obtained from the three fields and represented as a single data point on graphs. To determine significant differences among strains, a one-way ANOVA was performed on the mean values with strains in each experiment linked, followed by the Holm-Sidak test on pre-selected pairs of strains; GraphPad Prism v9 software was used for the graphs and analysis. *, p<0.05; **, p<0.01; ***, p<0.001.

### Cell lysates and immunoblots

Thirty OD_600_ units of cells were removed, centrifuged at 3000g for 5 min, and the pellets washed in 25 ml of H_2_O. Washed cell pellets were resuspended in 1 ml of H_2_O, transferred to microfuge tubes, and centrifuged at top speed in a microfuge for 1 min. Pellets were resuspended in 450 μl of H2O and chilled on ice. Fifty μl of 100% trichloroacetic acid was added, and the tubes vortexed and then allowed to remain on ice for 15 min. The suspension was centrifuged at 4 °C for 5 min and the supernatant was removed. Pellets were centrifuged for 15 sec, and the remaining supernatant was pipetted off. Three hundred μl of 2x Laemmli sample buffer was added, and the cells were resuspended by vortexing. To turn the suspension from yellow (from the residual acid) to blue, 4–5 μl of 5 N NaOH was added. Then, 250 mg of acid-washed glass beads were added, and the suspensions were subjected to three 1-min pulses in a mini-beadbeater (Biospec Products, Bartlesville, OK) in the cold room to lyse the cells. Samples were placed in a boiling water bath for 5 min and chilled. Leaving the beads behind, lysates were transferred to fresh tubes and centrifuged for 5 min at top speed. Supernatants were collected and used for immunoblotting.

The protein concentrations of the lysates were determined by an amido black filtration assay^56^ with Fraction V BSA (Sigma) for a standard curve. Then 20 μg of cell lysates were added to 10% SDS gels for polyacrylamide electrophoresis and detection of seipin; in parallel, 2 μg of lysates were run out for detection of G6PDH. Proteins were electroblotted from gels onto nitrocellulose. Blots were treated for 1 h or overnight with LI-COR PBS Blocking Buffer (diluted 1:4 in TBST) and then subjected to first and second antibody, with 3 5-min washes with TBST after each, before visualization on a LI-COR Biotechnology Odyssey Infrared Imaging System (Lincoln, NE) and quantification of bands using Image Studio v. 5.2.5 (LI-COR).

Antibodies for immunoblots: primary antibodies included anti-myc monoclonal 9E10 (Thermofisher, diluted 1:10,000) or anti-G6PDH (Sigma, diluted 1:20,000). Secondary antibodies included goat anti-mouse and goat anti-rabbit IRDye antibodies, used according to the manufacturer (LI-CORE).

---

**Table S1.**
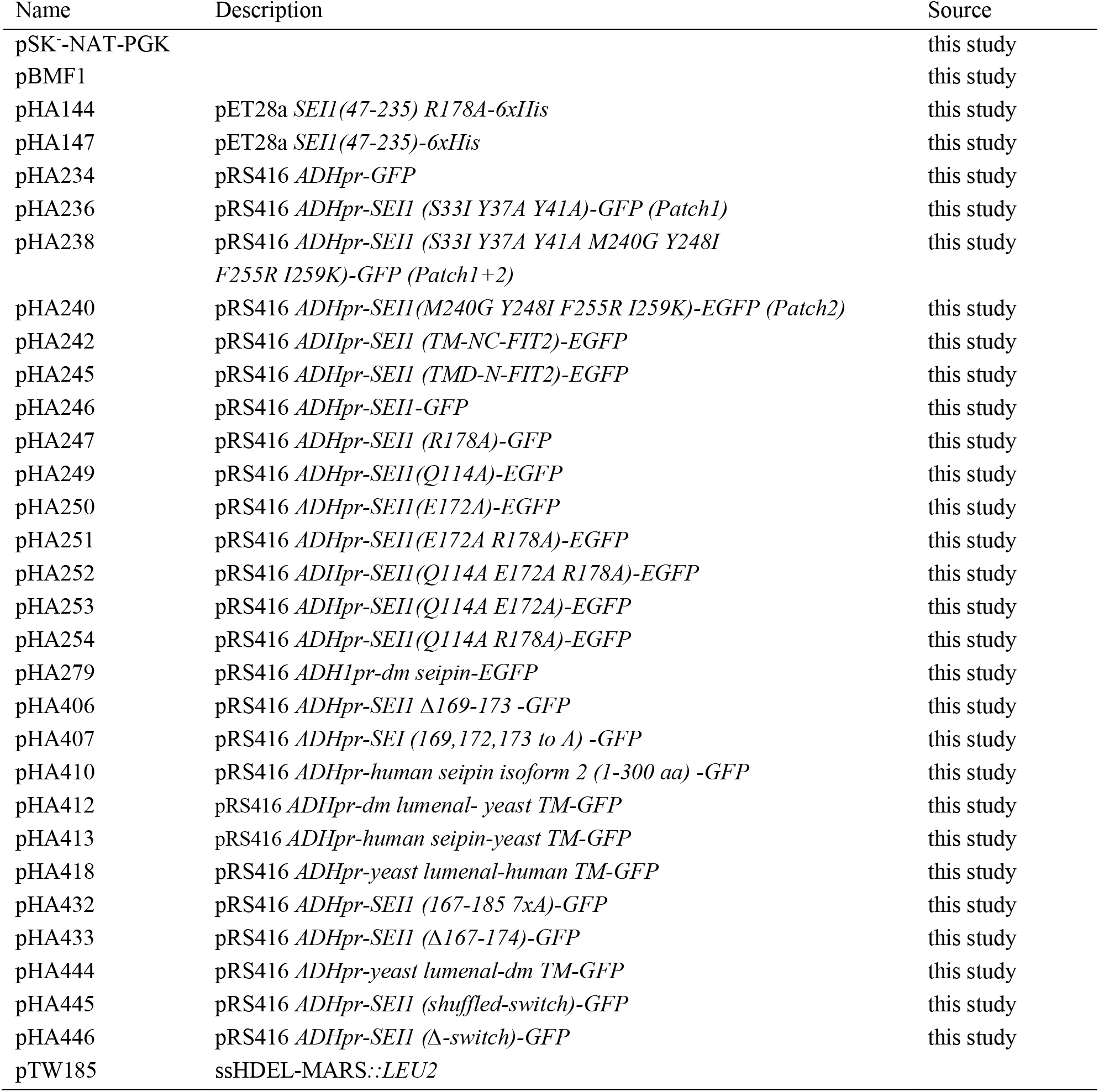
Plasmids used in this study.

**Table S2.**
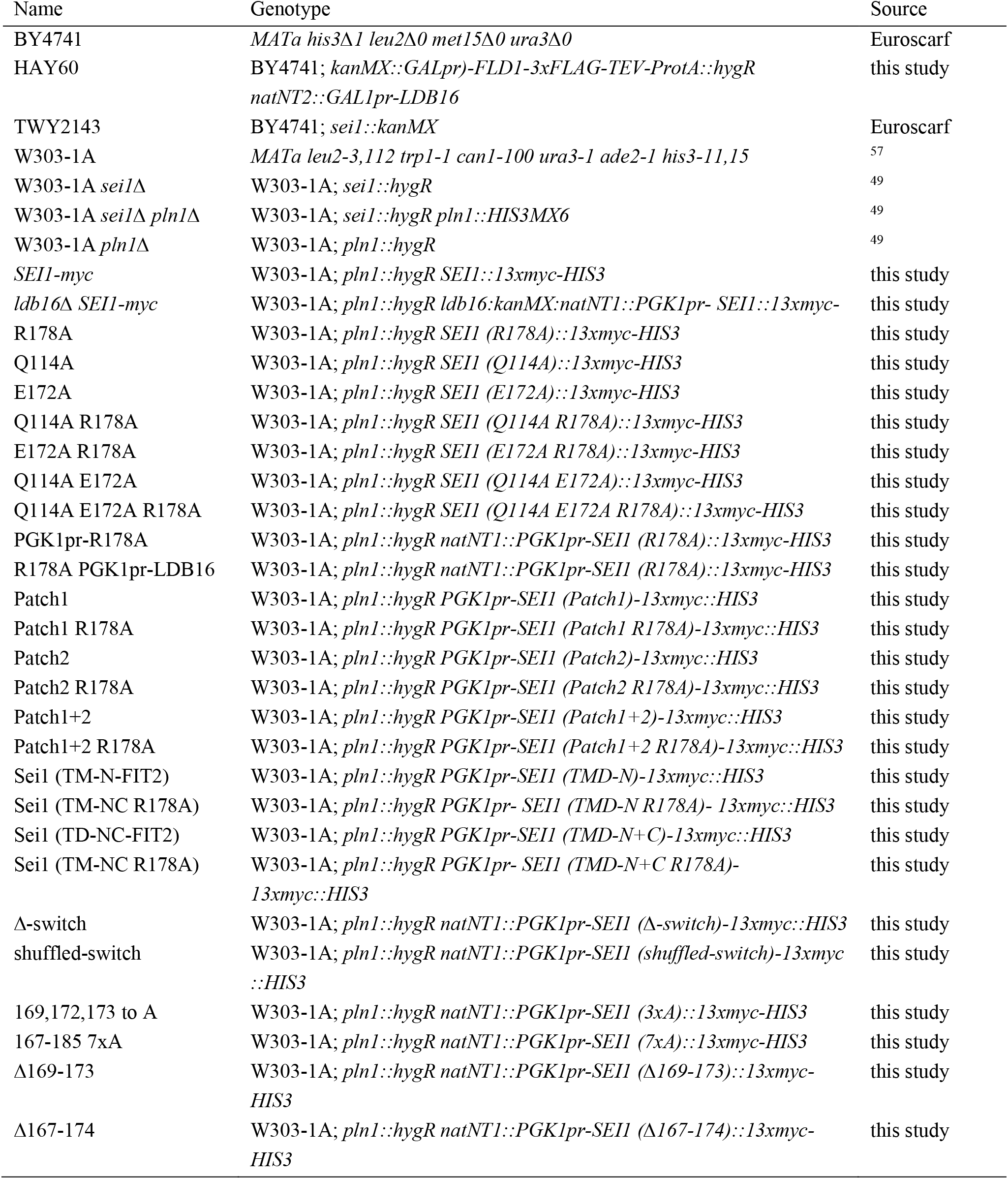
Yeast *S. cerevisiae* strains used in this study.

## Notes

### Competing Interest Statement

The authors have declared no competing interest.

